# Neuroblastoma immune microenvironment is characterized by limited T cell infiltration and abundant macrophages expressing targetable receptors

**DOI:** 10.64898/2026.11.09.764906

**Authors:** Ruth Nousiainen, Minttu Polso, Lassi Paavolainen, Aino Mutka, Veikka Valtanen, Satu Långström, Minna Koskenvuo, Virve Pentikäinen, Marjut Pihlajoki, Katja Eloranta, Vilja Pietiäinen

## Abstract

**Background & Aim:** Neuroblastoma (NBL) is a heterogenous pediatric malignancy ranging from low-risk infant tumors to high-mortality stage IV disease. We aimed to characterize the composition and spatial organization of the tumor immune microenvironment (TIME) across different tumor stages and assess the prevalence of immunotherapeutic targets, particularly focusing on macrophage-directed therapies.

**Methods:** Multiplexed immunofluorescence staining was performed on 185 retrospective formalin-fixed paraffin-embedded tissue cores from 39 NBL patients. Four antibody panels targeting a total of 32 proteins were used. Immune and stromal cell phenotypes and their spatial location were identified by single-cell image analysis and correlated with clinical features such as stage, MYCN status, prior treatment, and survival.

**Results:** Immune infiltration varied among samples and patients. CD45^+^ leukocytes comprised a median of 1.8% of all cells (range 0.2-51.0%, IQR 0.8-6.5%), dominated by tumor-associated macrophages (TAMs; 31.7% of CD45^+^ cells), and NK cells (15.6%). Stage IV tumors contained fewer lymphocytes. Chemotherapy shifted the TIME toward a macrophage-dominated, lymphocyte-depleted profile. Rather than segregating into M1- and M2-like subsets, TAMs co-expressed pro- and anti-inflammatory markers: CD40, CSF1R, and CD163 were broadly expressed, Clever-1 was detected on 41% of macrophages, and CD200R was virtually absent. CD47^+^CD45^−^ cells were enriched in stage IV tumors. In univariate Cox analysis, higher fractions of total CD45^+^, NK, NKT-like, B, and cytotoxic T cells were associated with improved overall survival. Higher proportion of CD47^+^CD45^-^ cells predicted worse survival. Blood leukocyte counts did not correlate with intratumoral immune infiltration.

**Conclusions:** The NBL TIME was heterogenous. High-risk tumors shared a lymphocyte-depleted, myeloid-skewed phenotype, whereas higher overall immune content predicted better survival. Macrophages were present in all tumors and expressed targetable receptors (CD40, CSF1R, CD163, Clever-1). CD47, enriched in high-risk disease, could also be a candidate macrophage-directed target.

## INTRODUCTION

Neuroblastoma (NBL), primarily arising in the sympathetic nervous system from neural crest progenitor cells, is the most common extracranial solid tumor in children ^1^. It is typically diagnosed in infancy with a global incidence of 0.28/100 000 in 2021 ^2,3^. Most cases are thought to be sporadic, although known NBL predisposing conditions include Beckwith-Wiedemann syndrome and Li-Fraumeni syndrome as well as germline mutations in the anaplastic lymphoma kinase (*ALK)*, paired-like homeobox 2B (*PHOX2B),* and kinesin family member 1B (*KIF1B)* genes. NBL is clinically heterogeneous, ranging from spontaneous regression of biologically favorable tumors in infants, to metastatic, high-risk disease associated with high mortality in older children. Despite treatment advances, NBL remains responsible for a disproportionate share of pediatric cancer mortality accounting for approximately 10% of all pediatric cancer deaths ^3–5^.

The role of tumor immune microenvironment (TIME) is thought to be critical in cancer development, progression, and therapy response. The NBL TIME has generally been considered immunosuppressive or immunologically cold ^6^. Higher B cell, NK cell, and T cell infiltration has been associated with improved prognosis in NBL, and localized tumors often contain well organized tertiary lymphoid structures (TLSs) ^7,8^. In contrast, high-risk NBLs show low tumor infiltration by T cells, NK cells, and B cells, and TLSs are found to be either immature or not present ^8,9^. Immunogenicity also differs between NBL tumor cell lineage states. Tumors with a mesenchymal gene expression profile show higher expression of immune activation genes and signatures of T and NK cell infiltration than adrenergic tumors ^10^. *In vivo*, mesenchymal cells have been more responsive to immune checkpoint blockade ^10^. Macrophages are often abundant in the TIME and, depending on their functional state, they may either inhibit or support tumor progression. The infiltration of tumor-promoting CSF1R-expressing myeloid cells, including macrophages, has been correlated with poor clinical outcome in NBL ^11^. A recent single-cell transcriptomics study demonstrated that in NBL, macrophages shift towards immunosuppressive phenotype following chemotherapy, suggesting that therapy-induced TIME rewiring may contribute to treatment resistance ^12^.

Tumors can be targeted by modifying the TIME, for example by preventing the immune escape of tumor cells. In NBL, however, immunotherapeutic approaches have been limited by low tumor immunogenicity, particularly in high-risk tumors. An important exception is the anti-GD2 therapy in high-risk NBL ^13^. GD2 is overexpressed in NBL and anti-GD2 antibodies, such as dinutuximab, bind to it, triggering antibody-dependent cellular cytotoxicity and complement-dependent cytotoxicity. Anti-GD2 therapy is now a standard part of maintenance therapy for high-risk NBL and has improved the outcomes of high-risk NBL ^14,15^. Macrophage-targeting immunotherapies, such as anti-CD47 or anti-CSF1R, are emerging as promising treatment strategies in other cancers but remain mostly unexplored in NBL ^11,16^.

Based on previous evidence linking immune infiltration with NBL outcome, we hypothesized that clinically aggressive NBLs are characterized by distinct immune microenvironmental features, including reduced lymphoid infiltration and altered spatial organization of immune cell populations, and that these features may be associated with patient outcome. We used multiplexed immunofluorescence (mIF) to characterize the composition and spatial organization of the NBL TIME and integrated these findings with clinical data. We further assessed potential immunotherapeutic targets, with particular emphasis on macrophage-directed therapies, to identify features of the tumor microenvironment that may inform future therapeutic strategies.

## RESULTS

### Patient and cohort characteristics

A total of 39 patients diagnosed with NBL between 1995-2018 in Helsinki University Hospital were included in the study. An overview of patient characteristics is shown in **Table 1**. Median age at diagnosis was 1.7 years (range 0-12 yrs), 49% were male, 62% were classified as high-risk disease, and 41% had a MYCN amplification in tumor. Retrospective formalin-fixed paraffin-embedded (FFPE) biobank samples from these patients were obtained from Helsinki Biobank. Given the multi-decade span of samples, patients were treated according to the standard-of-care protocol applicable at the time of diagnosis and thus, treatment was therefore not uniform across the cohort.

**Table 1.** Patient characteristics.

| **Characteristic** | **N = 39, N (%) or median (range)** |
|---|---|
| Age at diagnosis (years) | 1.7 (0-12) |
| Sex |  |
| Female | 20 (51%) |
| Male | 19 (49%) |
| International Neuroblastoma Staging System (INSS) |  |
| Stage I | 5 (13%) |
| Stage II | 4 (10%) |
| Stage III | 5 (13%) |
| Stage IV | 25 (64%) |
| International Neuroblastoma Risk Group (INRG) |  |
| Very low | 5 (13%) |
| Low | 6 (15%) |
| Intermediate | 2 (5%) |
| High | 24 (62%) |
| Unknown | 2 (5%) |
| MYCN status |  |
| Not amplified | 18 (46%) |
| Low amplification | 5 (13%) |
| High amplification | 11 (28%) |
| NA | 5 (13%) |
| Metastatic disease at diagnosis | 26 (67%) |
| Bone marrow involvement | 25 (64%) |
| Shimada classification |  |
| Favorable histology | 17 (44%) |
| Unfavorable histology | 16 (41%) |
| NA | 6 (15%) |
| Received neoadjuvant chemotherapy | 23 (61%) |
| Received adjuvant chemotherapy | 31 (86%) |
| Relapse | 14 (36%) |
| Deceased | 19 (49%) |
| Median time to death (months; n = 19) | 13.5 (0.6-156) |

### NBL tumors show heterogenous immune infiltration with a prominent macrophage population

Multiplexed immunofluorescence (mIF) staining was performed on FFPE NBL samples using 32 antibodies (**Table 3** in Methods, **Figure 1A-B**), generating single-cell data on key tumor microenvironment (TME) populations. mIF revealed considerable inter-patient heterogeneity in the immune cell composition of NBL tumors (**Figure 2A**). The majority of patients showed low overall intratumoral immune infiltration. CD45^+^ leukocytes comprised a median of 1.8% of all cells (range 0.2-51.0%, IQR 0.8-6.5). The CD45^+^ compartment was dominated by tumor-associated macrophages (TAMs; CD45^+^CD68^+^HLA-DR^+/–^; 31.7% of CD45^+^ cells), followed by NK cells (CD45^+^CD3^-^CD56^+^; 15.6%), although their relative proportions varied considerably between patients. B cells (CD45^+^CD20^+^), T cells (CD45^+^CD3^+^CD56□), and NKT-like cells (CD45^+^CD3^+^CD56^+^) were found to be present in most tumor tissues but showed marked inter-patient variation. Representative images of two tumor cores with relatively high and low immune cell infiltration within our cohort and their corresponding cell-level spatial phenotype maps are shown in **Figure 2C-D**.

**Figure 1.**
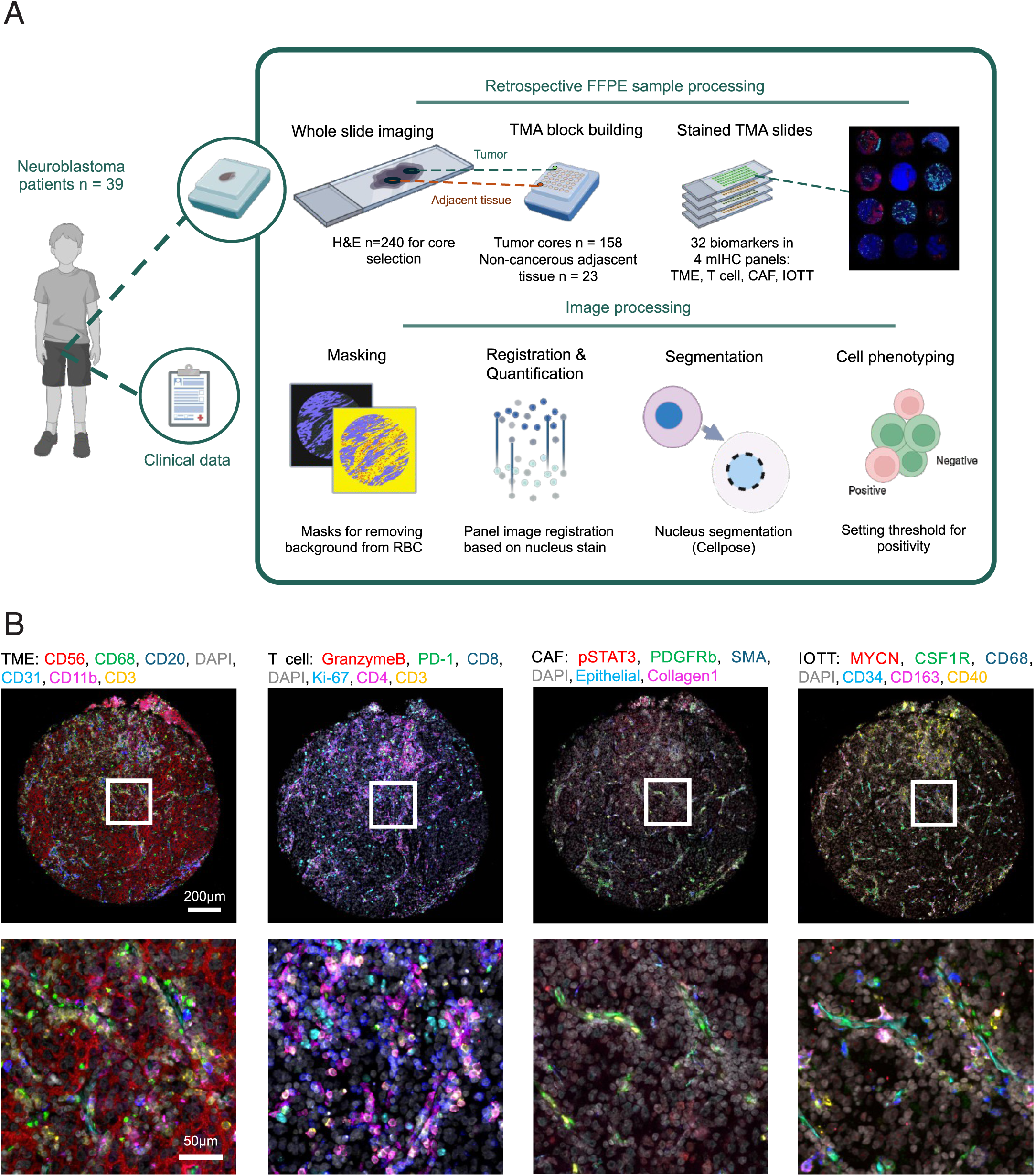
Overview of the study and mIF panels. **A**: Overview of study design. Clinical data were integrated with whole-slide H&E imaging, tissue microarray construction, and multiplex immunofluorescence analysis of retrospective FFPE NBL samples. Representative tumor regions were selected from whole-slide images for tissue microarray preparation, followed by staining with multiple immune, proliferation, and tumor-associated markers. Digitized tissue images underwent image preprocessing, including masking of red blood cells to exclude autofluorescence, registration of staining rounds, cell segmentation, and feature extraction. Quantitative image analysis was subsequently performed to characterize cellular composition, marker expression patterns, and spatial relationships within the tumor microenvironment. Created with BioRender.com. **B**: Representative multiplex immunofluorescence images showing selected markers from each of the four staining panels (TME, T cell, CAF, and IOTT). Scale bars: 200 µm in TMA core images and 50 µm in magnified regions.

**Figure 2.**
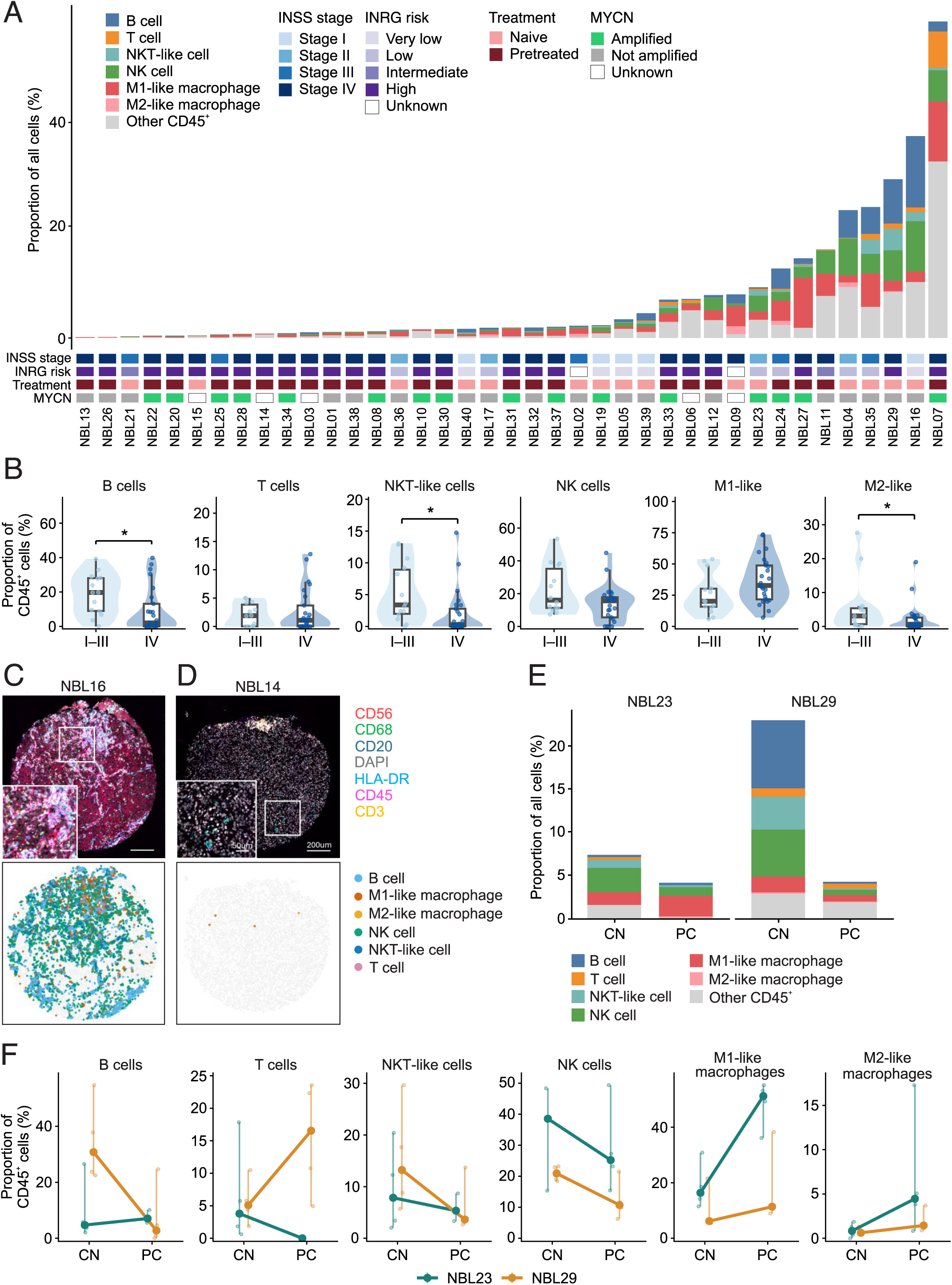
Overall immune composition of NBL tumor microenvironment and effects of chemotherapy on TIME composition. **A**: Proportion of each main immune cell type out of total cells in NBL tumor tissues. Bars represent patient-level median values. For NBL23 and NBL29, chemotherapy-naïve samples are shown. **B**: Comparison of INSS stage I-III (n = 14) vs stage IV (n = 25) tumors. Each dot represents the median of tumor cores from each patient. Boxplot shows the median and quartile values with whiskers extending to the most extreme non-outlier values (1.5 × IQR). P-values were calculated using Mann-Whitney U test with Benjamini-Hochberg FDR correction. * = p-value < 0.05. **C-D**: Representative mIF images and corresponding spatial cell maps of tumors with relatively high (NBL16; **C**) and relatively low (NBL14; **D**) total immune cell infiltration. Scale bar 200 μm; close-up image 50 μm. **E**: Comparison of total immune cell proportion out of all cells in two patients with paired samples (NBL23 and NBL29) before (CN) and after (PC) neoadjuvant chemotherapy. Medians of tumor cores are shown. **F**: Change in the proportions of immune cell populations in the paired longitudinal samples, NBL23 and NBL29. Small dots = individual cores. Large dots = median. Vertical bars = range across cores.

### Effects of disease stage, risk group, and prior chemotherapy on overall immune infiltration

Immune cell composition was compared between stage IV and I-III tumors (**Figure 2B**). B cells, NKT-like cells, and M2-like macrophages were less abundant in stage IV tumors. The median B cell fraction of total CD45^+^ cells was 2.0% in stage IV vs 19.7% in stage I-III (p = 0.044), NKT-like cells 0.3% vs 3.4% (p = 0.042), and M2-like macrophages 0.6% vs 3.1% (p = 0.044). M1-like macrophages tended to be more prevalent in stage IV tumors (32.8% vs 20.2%) but the difference was not significant after correction for multiple testing (p = 0.067). T cell and NK cell proportions did not differ significantly between the groups. Likewise, the immune composition differed by the International neuroblastoma risk group (INRG) classification (**Supplementary Figure S1A**). High-risk tumors were overall less immune infiltrated (median total CD45^+^ 1.17% of all cells vs 4.48% in lower-risk tumors, p = 0.049). Compared to very low, low, and intermediate risk tumors, high-risk tumors contained lower proportions of NKT-like cells (3.6% vs 0.4%, p = 0.030) and NK cells (24.5% vs 14.1%, p = 0.030), and a higher proportion of M1-like macrophages (20.0% vs 33.1%, p = 0.030). B cells were also lower in high-risk tumors (19.8% vs 2.8%), although this did not reach significance after correction (p = 0.071).

TIME composition differed between chemotherapy-naïve and post-chemotherapy samples (**Supplementary Figure S1B**). It is good to note that treatment status was largely confounded with stage, as most stage IV tumors had received neoadjuvant chemotherapy prior to sampling. Compared with chemotherapy-naïve (CN) samples, samples that were obtained post-neoadjuvant chemotherapy (PC) contained lower proportions of B cells (19.6% vs 3.1%, p = 0.027), NKT-like cells (4.4% vs 0.8%, p = 0.027), NK cells (20.2% vs 12.0%, p = 0.040), and M2-like macrophages (4.9% vs 0.7%, p = 0.027), whereas M1-like macrophages (HLA-DR^+^) were markedly more prevalent in PC samples (20.0% vs 42.1%, p = 0.027). The proportion of T cells did not differ between the groups. Overall immune infiltration, measured as proportion of CD45^+^ cells of all cells, tended to be higher in CN tissues (2.7% vs 1.5%), but this difference was not statistically significant (p = 0.086).

Two NBL patients had longitudinal paired samples, collected before and after chemotherapy. NBL23 (< 12 months at diagnosis, INSS stage II/L2/low risk) underwent primary resection without chemotherapy (first sample, chemotherapy-naïve = CN), was assigned to follow-up only. Nine months later, the patient had a metastatic relapse and received neoadjuvant chemotherapy followed by resection of the relapsed tumor (second sample, post-chemotherapy = PC). NBL29 (age group 2-7 years at diagnosis, INSS stage IV/M/high risk) was sampled at diagnosis before any treatment (first sample, CN) and another sample was obtained after induction chemotherapy at tumor resection (second sample, PC). Both patients received induction therapy combining vincristine, etoposide, cisplatin, cyclophosphamide, and doxorubicin, and neither received radiotherapy before resection. Both post-chemotherapy samples contained viable tumor, and thus both patients subsequently received high-dose chemotherapy, autologous stem cell transplantation and local radiotherapy (no samples collected). NBL23 was alive at last follow-up more than 10 years after diagnosis whereas NBL29 was deceased (OS 48 months). In both longitudinal sample pairs, overall CD45^+^ immune infiltration decreased after chemotherapy, from 7.2% to 4.2% of all cells in NBL23 and from 23.0% to 4.3% in NBL29 (**Figure 2E**). Within the CD45^+^ compartment, total macrophages increased after treatment in both patients (17.2% to 56.7% and 6.8% to 12.8%) and NKT-like cells decreased (7.8% to 5.3% and 13.2% to 3.6%) (**Figure 2F**). B cells and T cells showed mixed responses: B cells increased in NBL23 (4.7% to 7.1%) but decreased in NBL29 (30.8% to 2.8%), while T cells decreased in NBL23 (3.8% to 0.0%) and increased in NBL29 (5.1% to 16.5%). Because only two patients contributed paired samples, no statistical testing was performed on these comparisons.

The CD56^+^CD45^−^CD31^−^PDGFRβ^−^ tumor-associated cell compartment showed marked inter-patient heterogeneity. At patient level, the compartment comprised a median of 57.2% of all cells (IQR 31.1-73.3%), with values ranging from 0.2% to 87.9% (**Figure 3A**). Five of 39 tumors had a median compartment proportion below 10%, and nine had values below 25%. The proportion of the CD56^+^ tumor-associated compartment was lower in stage IV than in stage I-III tumors (**Figure 3B**). These findings indicate substantial variation in the representation of the CD56^+^ tumor-associated compartment, with lower levels associated with more advanced disease.

**Figure 3.**
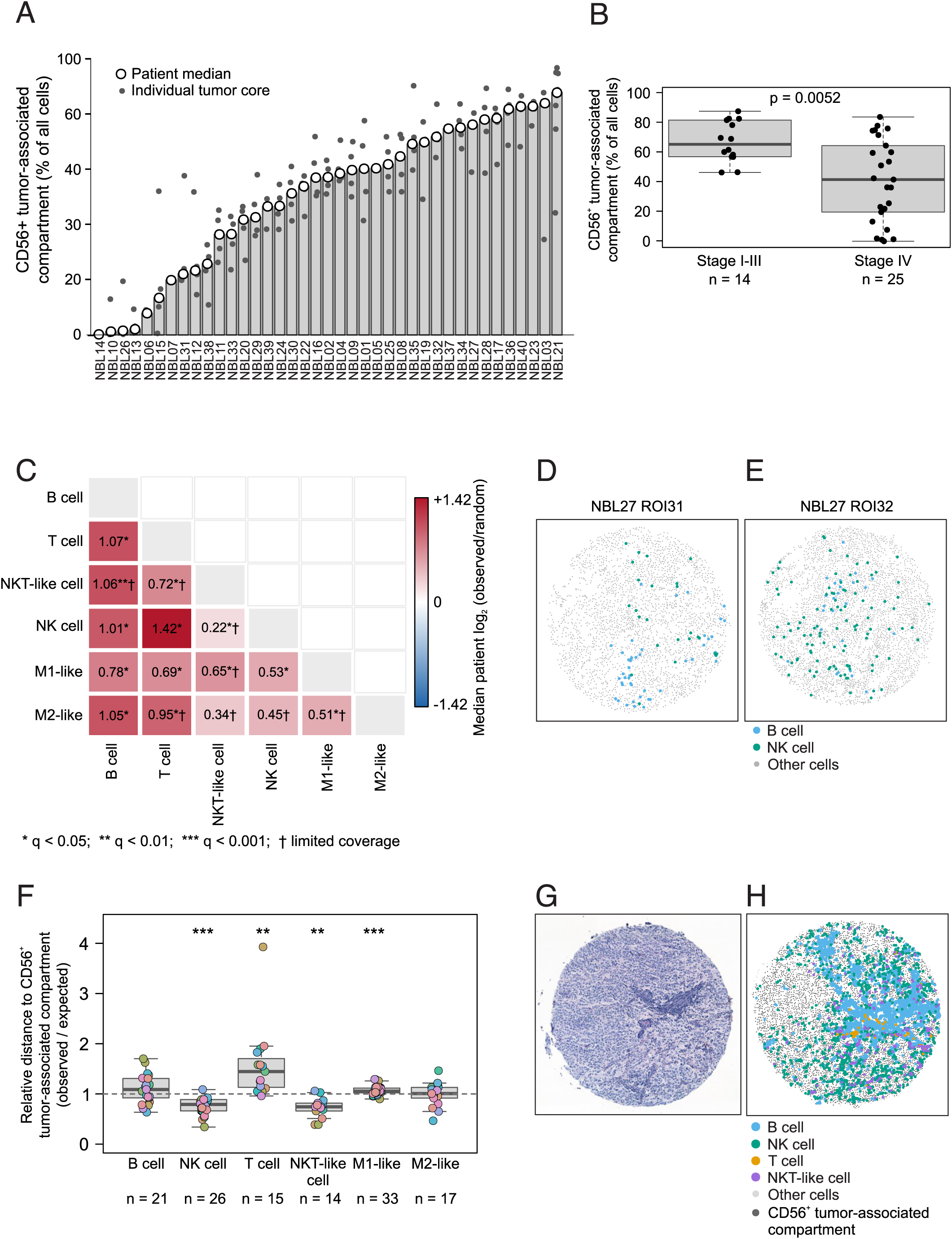
Spatial immune landscape of NBLs. **A**: CD56^+^CD45^−^CD31^−^PDGFRβ^−^ tumor-associated compartment across NBLs. Bars represent the median proportion (open circles) across tumor cores for each patient (n = 39), ordered from lowest to highest; gray dots indicate individual tumor cores. For patients with paired pre- and post-treatment samples (NBL23 and NBL29), only pretreatment tumor cores were included. **B**: Proportion of the CD56^+^CD45^−^CD31^−^PDGFRβ^−^ tumor-associated compartment in stage I-III (n = 14) vs stage IV (n = 25) tumors. Each dot represents the median of one patient. **C**: Heatmap showing patient-level median log2-transformed observed-to-random nearest-neighbor ratios for all pairwise comparisons among immune-cell populations. Positive values (red) indicate spatial separation relative to the within-core random-label expectation; negative values (blue) would indicate relative spatial proximity. Benjamini-Hochberg correction across all 15 comparisons (q < 0.05). † = comparisons with limited coverage, defined as <8 evaluable patients or <15 evaluable TMA cores. **D-E**: Spatial maps from two cores of NBL27 illustrating intratumoral heterogeneity in B–NK organization. ROI31 showed clear B–NK spatial separation (observed/random ratio 1.62, empirical P = 0.001), whereas ROI32 showed little evidence of separation (ratio 1.09, P = 0.110). **F**: Localization of the main immune cell populations relative to the CD56^+^CD45^−^CD31^−^PDGFRβ^−^ tumor-associated compartment (Gate C) across evaluable NBL tumor cores. Ratios <1 indicate relatively Gate C-proximal localization, whereas ratios >1 indicate relatively Gate C-distal localization. Core-level ratios were summarized as the median across evaluable tumor cores for each patient. Colored points represent patient-level median ratios. Each color represents one patient. Only cores containing at least 10 Gate C cells and immune cells of interest were included. Two-sided Wilcoxon signed-rank tests against a ratio of 1, BH correction applied. **G-H**: Representative H&E and spatial maps of NBL16 ROI65 illustrating the tumor-like compartment and lymphoid-cell localization. ROI = region of interest, TMA core number.

### Immune cell populations show widespread spatial separation in immune rich NBL tumors

Given the marked heterogeneity in immune infiltration, we next examined the spatial organization of immune cells in immune-rich tumors (defined as the upper quartile of patient-level median CD45^+^ infiltration; 10 of 39 patients; 36 malignant cores). For each pair of six mutually exclusive phenotypes (B, T, NK, NKT-like cells and M1-like HLA-DR^+^ and M2-like HLA-DR^−^ macrophages), observed cross-nearest-neighbor distances were compared with those expected under random relabeling of cells. Median patient-level observed-to-random distance ratios exceeded 1 for all 15 pairs, indicating spatial separation rather than random intermixing (**Figure 3C**). After Benjamini-Hochberg correction, separation was significant for T–NK, B–T, B–NK, B–M1-like, T–M1-like, and NK–M1-like pairs. Comparisons involving NKT-like or M2-like cells showed similar trends but included fewer evaluable tumor cores. Spatial separation metrics were not associated with INSS stage, age at diagnosis, or MYCN status after multiple-testing correction.

B and NK cells were further examined because both populations were well represented and clearly separated in the pairwise analysis (**Supplementary Figure S2A**). Of the 36 immune-rich tumor cores from 10 patients, 29 contained ≥10 B cells and ≥10 NK cells. All 29 evaluable cores had B–NK observed-to-random distance ratios above 1 (core-level median 2.02), and the spatial separation was significant in 27 cores after multiple-testing correction. At the patient level, the median observed-to-random ratio was 2.20 and exceeded 1 in all 10 patients (Wilcoxon signed-rank p = 0.0059). Despite this consistent overall B–NK separation, spatial organization varied between the tumor cores from the same patient. In NBL27, core ROI31 showed clear B–NK separation (ratio 1.62, permutation p = 0.001), whereas ROI32 was close to random organization (ratio 1.09, p = 0.110; **Figure 3D-E**). Thus, although B and NK cells generally occupied distinct spatial niches in immune-rich tumors, the extent of separation varied locally within an individual tumor.

To further explore if immune-cell patterns varied across tissue regions, tumor cores were compared with tumor-border and adjacent non-cancerous tissues. Tumor-border cores were available from only four patients and were therefore analyzed descriptively. B cell infiltration generally decreased from tumor toward tumor-border, whereas NK cell and M2-like macrophage patterns were more heterogeneous. A representative transition from tumor core to tumor-border in patient NBL33 is shown in the **Supplementary Figure S2B-E**.

### Immune cell populations show distinct localization relative to the CD56^+^ tumor-associated compartment

We next assessed the localization of immune cells relative to the CD56^+^CD45^−^CD31^−^PDGFRβ^−^ tumor-associated cells (hereafter called CD56^+^ tumor cells). For each immune cell type, the mean distance to the nearest CD56^+^ tumor cell was divided by the corresponding mean for all six immune populations in the same core. Ratios <1 thus indicate tumor-proximal localization and ratios >1 tumor-distal localization. Localization relative to CD56+ tumor cells differed between immune populations (**Figure 3F**). NK and NKT-like cells were mostly CD56^+^ tumor-proximal (median patient-level ratios 0.79 and 0.74, FDR-adjusted p < 0.001 and 0.0031). In contrast, T cells were consistently more distal from the CD56^+^ tumor cells (median ratio 1.45; p-adj = 0.0018). M1-like macrophages showed a modest distal shift (median ratio 1.06; p-adj < 0.001). The localization of B cells and M2-like macrophages relative to the CD56^+^ tumor cells showed no consistent pattern. None of the localization patterns of these immune populations was associated with INSS stage, age at diagnosis or MYCN status after multiple-testing correction. NBL16 ROI65 is shown as an example of the distribution of immune-cell populations relative to the CD56^+^ tumor-associated compartment (**Figure 3G-H**). These findings demonstrate cell-type-specific spatial organization relative to the tumor-associated compartment, with NK and NKT-like cells preferentially localized closer to CD56^+^ tumor cells and T cells showing more distal localization.

### Cytotoxic T cells express exhaustion markers across risk groups in NBL

In most NBL tumor cores, the overall T cell infiltration was low. The majority of cytotoxic T cells (CD3^+^CD8^+^) exhibited markers of functional inactivity: 58% were PD-1^+^ and 79% TIM-3^+^ (patient-level medians; **Figure 4A**). Only 10% of CD3^+^CD8^+^ T cells expressed Granzyme B, a marker of cytotoxic capacity (**Figure 4A-B**). The co-expression of inhibitory receptors was extensive: 97% of PD-1^+^CD8^+^ T cells were also TIM-3^+^, accounting for a median of 55% of all CD8^+^ T cells (**Figure 4C**). Nearly all of the few Granzyme B□ CD8□ T cells also expressed TIM-3, indicating that TIM-3 expression was not restricted to cells lacking cytotoxic capacity. Regulatory T cells (CD3^+^CD8^−^CD4^+^FOXP3^+^) represented a minor population, comprising a median of 1.7% of all CD3^+^ T cells. None of the T cell subpopulations differed significantly between stage I-III and stage IV tumors, and inter-patient variability was considerable for every subpopulation (**Figure 4A**).

**Figure 4.**
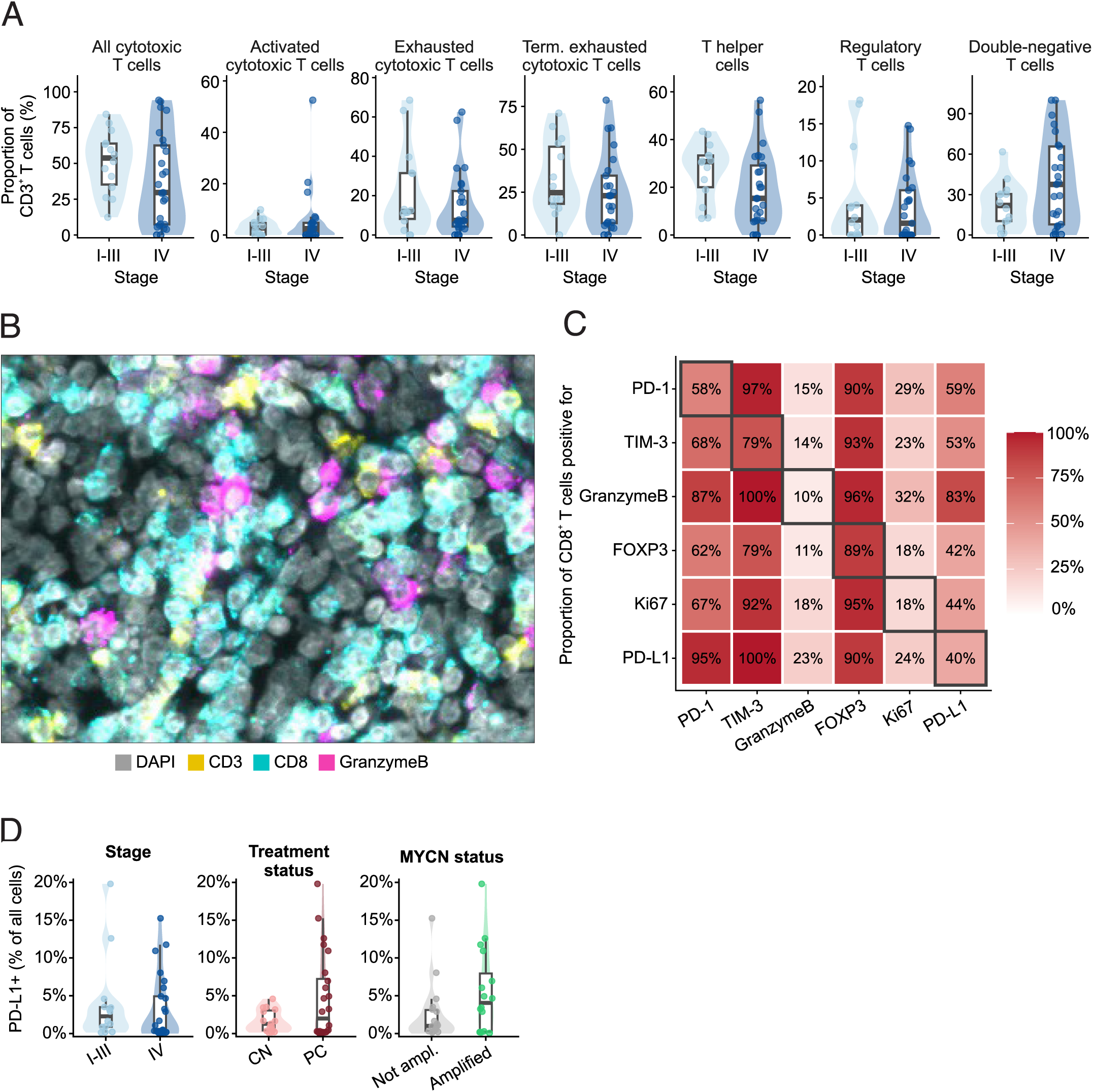
T cells widely exhibit exhaustion markers in NBL tumor tissues. **A**: Violin plots show the proportions of different T cell subpopulations as a percentage of total CD3^+^ cells. These consist of cytotoxic T cells (CD3^+^CD8^+^), activated cytotoxic T cells (CD3^+^CD8^+^GranzymeB^+^), exhausted cytotoxic T cells (CD3^+^CD8^+^PD-1^+^), terminally exhausted cytotoxic T cells (CD3^+^CD8^+^TIM-3^+^), T helper cells (CD3^+^CD8^−^CD4^+^), regulatory T cells (CD3^+^CD8^−^CD4^+^FOXP3^+^), and double-negative T cells (CD3^+^CD8^−^CD4^−^). Each dot represents the median of one patient. Boxplot shows the median and quartile values with whiskers extending to the most extreme non-outlier values (1.5 × IQR). Statistical comparisons were performed using the Mann-Whitney U test with BH correction; none reached significance. For patients with paired samples (NBL23 and NBL29), only the chemotherapy-naïve sample is shown. **B**: Representative multiplex immunofluorescence image of the T-cell compartment: CD3 (yellow), CD8 (cyan), Granzyme B (magenta) and nuclei (DAPI, grey). **C**: Co-expression of T cell markers within the CD8^+^ T-cell compartment. Cell counts were pooled across each patient’s tumor cores containing ≥10 CD8^+^ cells (60 of 157 cores, 23 of 39 patients), a fraction was calculated per patient, and squares show the median across patients. **D**: PD-L1 expression compared between stage I-III vs stage IV tumors, chemotherapy-naïve (CN) vs post-neoadjuvant chemotherapy (PC) samples and between MYCN-amplified and non-amplified tumors.

PD-L1 was detectable in nearly every tumor but at low levels with patient-level median expression being 1.28% of all cells (IQR 0.21-4.63%). 74% of PD-L1^+^ cells were CD3^−^. Because the T cell panel lacked an NBL-specific tumor marker, PD-L1 expression could not be assessed specifically on tumor cells. Overall PD-L1 expression was lower in stage IV tumors, but the result was not statistically significant (**Figure 4D**).

### Tumor-associated macrophages in NBL frequently co-express pro-inflammatory and anti-inflammatory markers

Tumor-associated macrophage (TAM) phenotypes varied considerably between tumors. CD45^+^CD68^+^HLA-DR^+^ (M1-like) TAMs were substantially more frequent than HLA-DR^−^(M2-like) TAMs, comprising patient-level medians of 29.5% and 0.85% of CD45□ cells (0.54% and 0.03% of all cells), respectively, with substantial heterogeneity between patients (**Figure 5A**). Next, macrophage-associated therapeutic targets were further characterized (**Figure 5B-C**). CD40 (a costimulatory receptor associated with pro-inflammatory, M1-like polarization), CSF1R (a receptor driving macrophage survival associated with anti-inflammatory, M2-like polarization) and CD163 (a scavenger receptor associated with M2-like polarization) were broadly expressed, with patient-level median positivity of 99.9% (IQR 94.9-100), 96.6% (IQR 91.8-100), and 90.3% (IQR 82.6-94.4) of all CD45^+^CD68^+^ macrophages respectively. Clever-1 (stabilin-1; a scavenger receptor associated with M2-like polarization) expression was more heterogenous (median 40.9%, IQR 29.8-64.6). In contrast, CD200R^+^ macrophages (an inhibitory receptor) were detected in low numbers in only two of 39 patients. Although 19 of the 32 possible marker combinations occurred, co-expression was concentrated in two phenotypes (**Figure 5D**). CD163^+^CD40^+^CSF1R^+^Clever^−^ (median 38.1% of CD45^+^CD68^+^ macrophages, IQR 19.7-51.9) and CD163^+^CD40^+^CSF1R^+^Clever^+^ (33.9%, IQR 26.5-58.8). Together, these two phenotypes made up a median of 79.2% of all CD45^+^CD68^+^ macrophages. The next most common combination was CD45^+^CD68^+^CD40^+^CSF1R^+^ (6.4%). All remaining combinations were rare, each averaging under 4%. Thus, most NBL TAMs express a mixed activation phenotype rather than discrete M1/M2 polarization, many carry potentially targetable immunosuppressive receptors.

**Figure 5.**
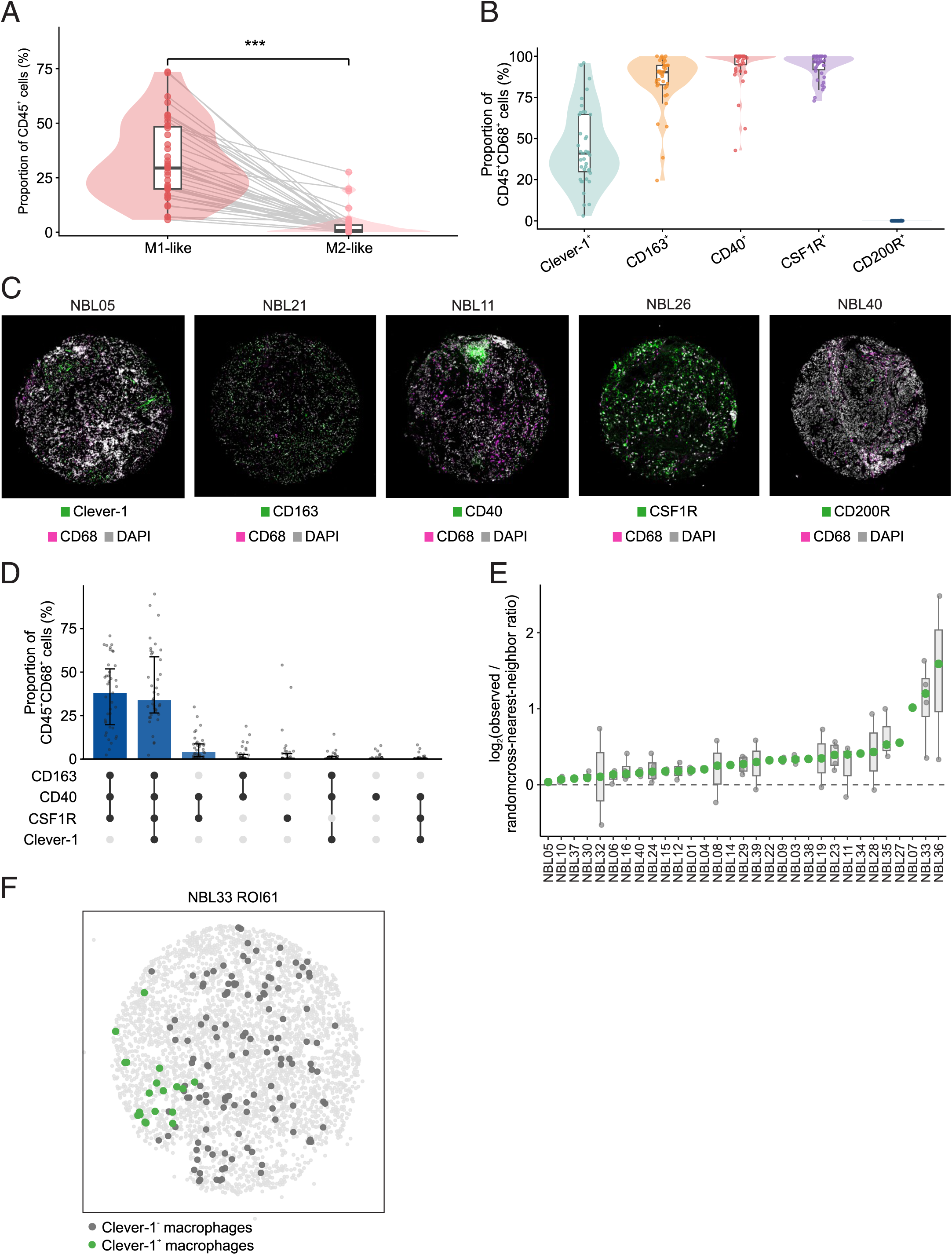
Tumor associated macrophages in NBL co-express M1- and M2-associated markers rather than segregating into polarized phenotypes. **A**: M1-like (CD45^+^CD68^+^HLA-DR^+^) vs M2-like (CD45^+^CD68^+^HLA-DR^-^) macrophages as a proportion of total CD45^+^ cells. Data from TME panel. Lines join each patient. Paired Wilcoxon signed-rank, *** p < 0.001. **B**: Proportion of CD45^+^CD68^+^ macrophages positive for each macrophage marker (Clever-1, CD163, CD40, CSF1R, CD200R) regardless of the expression of the other markers. Single cell data from IOTT panel. Each dot represents the median of one patient. **C**: Representative images of CD68 (magenta) with each macrophage marker (green) and nuclei (grey). Each TMA core is 1 mm in diameter. **D**: UpSet plot of the top eight marker combinations among CD45^+^CD68^+^ macrophages. Each column is one complete phenotype, positive for the markers with filled dots and negative for those with empty dots. CD200R is not shown because it occurs in none of the top eight combinations. Bars show medians, whiskers IQR. Each dot is the median of one patient. **E**: Spatial segregation of Clever-1-positive and Clever-1-negative CD45^+^CD68^+^ macrophages. Small grey points = individual tumor cores, green points = patient medians (31 patients, 1-4 cores per patient). Values are shown as log2-transformed observed-to-random cross-nearest-neighbor ratios; values above 0 indicate greater separation than expected under within-core random-label permutation. **F**: Spatial map of NBL33 ROI61, showing pronounced segregation of Clever-1-positive and Clever-1-negative macrophages (observed/random ratio 3.09; log2 ratio 1.63; empirical p = 0.001; FDR = 0.0042).

Because Clever-1 expression distinguished the two dominant macrophage populations, we next examined whether the localization of Clever-1^+^ and Clever-1^−^ macrophages differed. The two populations were spatially segregated relative to random-label expectation (**Figure 5E-F**). The patient-level median observed-to-random cross-nearest-neighbor ratio was 1.21 (IQR 1.13-1.32) and exceeded 1 in all 31 evaluable patients (Wilcoxon p = 1.23 × 10□□). At the core level, 65 of 71 evaluable cores had ratios above 1, of which 27 cores remained significant after multiple-testing correction. Clever-1^+^ macrophages also showed modest same-phenotype clustering (median ratio 0.95, IQR 0.86–1.01; 21/31 patients <1; Wilcoxon p = 0.041). These findings suggest that Clever-1 expression identifies a spatially organized macrophage subpopulation within NBL tumors.

### CD47 expression varies in NBL tumor tissue

Non-leukocyte CD47^+^ cells (an anti-phagocytic ligand of the macrophage receptor SIRPα, mostly expressed on tumor cells, defined here as CD47^+^CD45^−^) were detected in all NBL tumors in this cohort. However, variation between patients was considerable (**Figure 6A**). The median fraction of CD47^+^CD45^−^ cells of all cells was 0.48%, ranging from 0.001% to 25.3%. Expression was higher in INSS stage IV than in stage I-III tumors (median 1.42% vs 0.13%, p = 0.0005) and in post-chemotherapy compared with chemotherapy-naïve samples (1.54% vs 0.18%, p = 0.0002; **Figure 6B**). The fraction of CD47^+^CD45^-^ cells was numerically higher in MYCN-amplified than in non-amplified tumors (1.54% vs 0.38%, n = 16 vs 18), but the difference did not reach significance (p = 0.09; **Figure 6B**). Among tumors resected after neoadjuvant chemotherapy, CD47^+^CD45^−^ proportion was unrelated to histological response: the median fraction was 1.67% in tumors with a good response (≤10% viable tumor, n = 11) and 1.88% in those with a partial or poor response (n = 12; p = 0.98; **Figure 6B**). Examples of the variability of CD47^+^ cell proportions are shown in **Figure 6C**: the near-absent staining in NBL05 (0.05% of all cells) and the abundant staining in NBL38 (14.0%).

**Figure 6.**
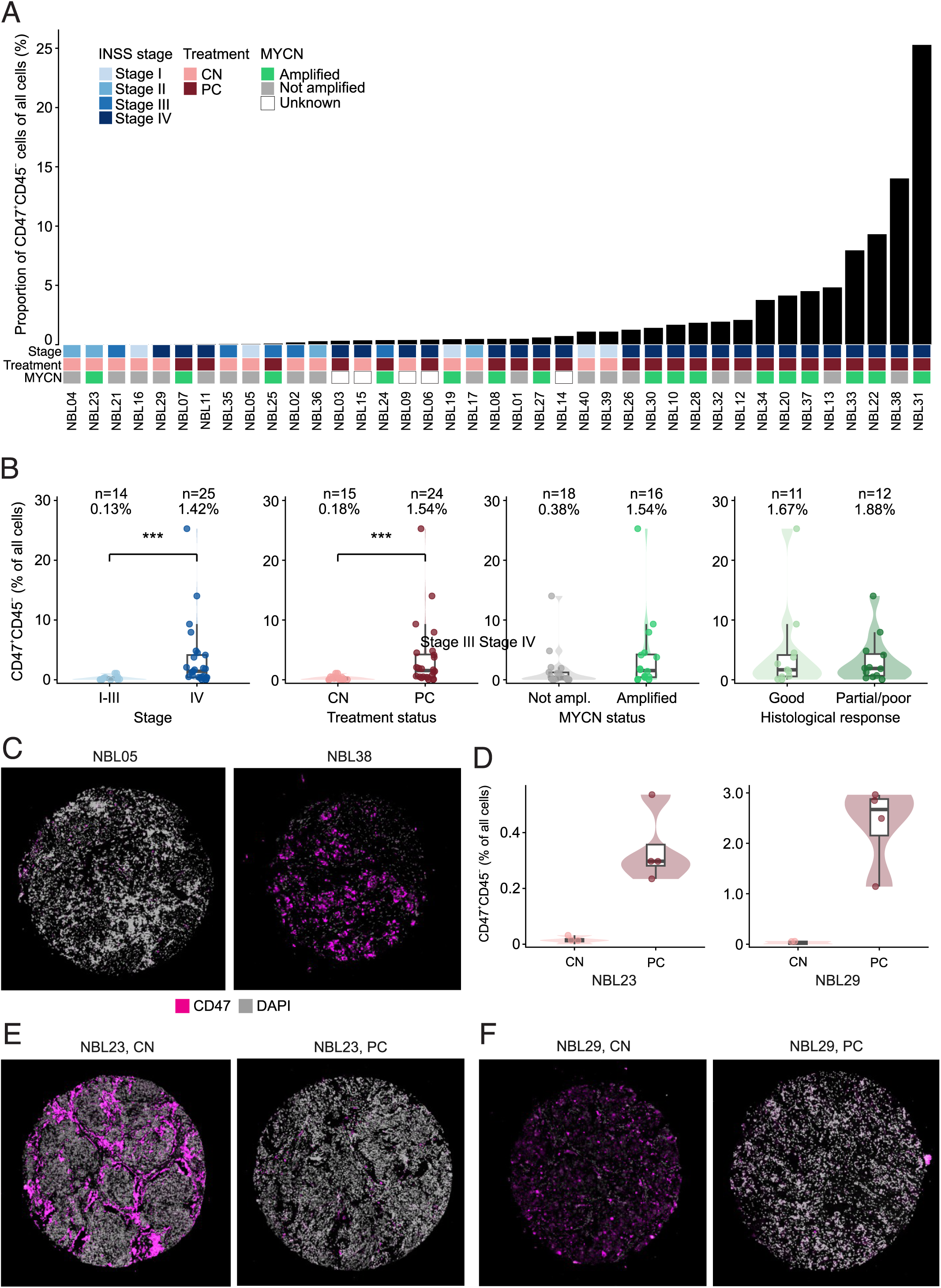
Higher CD47 expression is observed in higher-risk NBLs. **A**: The proportion of CD47^+^CD45^-^ cells of all cells in NBL tumor tissues, ranked from lowest to highest proportion. Each bar represents one patient (n = 39); values are patient-level medians. For paired-sample patients NBL23 and NBL29, only chemotherapy-naïve samples are shown. **B**: Proportion of CD47^+^CD45^-^ cells of total cells in stage I-III vs stage IV tumors, chemo-naïve (CN) vs post-neoadjuvant chemotherapy (PC), MYCN amplified vs non-amplified tumors and, in tumors resected after neoadjuvant chemotherapy, by histological response (good, ≤10% viable tumor, n = 11, vs partial/poor, n = 12). Histological response panel is restricted to post-chemotherapy samples (n = 23), in which NBL23 and NBL29 are represented by their post-chemotherapy sample. Each dot represents one patient. Mann-Whitney U; *** p < 0.001. **C**: Example images of high and low CD47-expressing tumor cores. Low CD47: NBL05 (ROI 19), high CD47: NBL38 (ROI 83). ROI = region of interest. **D**: CD47 expression in paired NBL samples NBL23 and NBL29 before and after chemotherapy. Each dot represents one punch. Inner boxplots show the median and interquartile range. **E-F**: Example images of the two paired samples before and after chemotherapy, NBL23 (**E**) and NBL29 (**F**). Red blood cell masks were applied during image analysis (not shown); note that part of the CD47 signal is vascular and appears more abundant in the images than the quantified cell fractions indicate. TMA cores are 1mm in diameter.

An increase in CD47^+^CD45^-^ cells was seen in both patients with paired samples obtained before and after chemotherapy. The median fraction of CD47^+^CD45^-^ cells rose from 0.01% to 0.30% in NBL23 and from 0.03% to 2.67% in NBL29 (**Figure 6D**); in both patients, CD47^+^CD45^-^ cells were close to absent before treatment. In the representative images, part of the CD47 signal originates from red blood cells, so CD47-positivity appears more abundant than the quantified cell fractions indicate. Red blood cells were masked during image analysis and excluded from quantification. (**Figure 6E-F**).

### Stromal composition and vascular content are heterogeneous and unrelated to outcome

Stromal markers were detected in all tumors with wide inter-patient variation (**Supplementary Figure S3A**). Among the stromal cell markers, PDGFRβ^+^ fibroblasts/pericytes were the most abundant (median 17.4% of all cells), followed by α-SMA^+^ myofibroblastic cells (11.2%), dual-marker PDGFRβ^+^α-SMA^+^ cancer-associated fibroblasts (CAFs) (8.1%), and pSTAT3^+^α-SMA^+^ myofibroblasts (7.8%). FAP^+^ CAFs were rare (0.69%). Collagen I staining was frequently observed in the extracellular matrix of NBL tumors, often showing a fibrillar or reticular pattern. The extent of staining varied considerably between samples, ranging from nearly absent staining to extensive staining in stroma-rich tissues. In our cell-based analysis, Collagen I-associated staining was detected in a median of 41.5% of segmented cells. None of the six stromal measures differed by stage, MYCN status, or INPC histology (**Supplementary Figure S3B**). Stromal content showed no association with outcome (**Supplementary Figure S3C**). For the dual-marker PDGFRβ^+^α-SMA^+^ CAF gate the hazard ratios for high versus low content (median split) were 0.58 (95% CI 0.23-1.47, p = 0.25) for overall and 0.75 (0.32-1.77, p = 0.52) for event-free survival.

Tumor vascular content, estimated as the proportion of CD34^+^CD45^-^Epithelial^−^ cells, varied between patients (median 2.97% of all cells, IQR 2.12-5.39%; **Supplementary Figure S4A**). It was numerically higher in stage IV than in stage I-III tumors (3.59% vs 2.32%, p = 0.072) and in MYCN-amplified than in non-amplified tumors (3.46% vs 2.69%, p = 0.29), but neither difference reached significance. Vascular content was likewise unrelated to overall survival in our cohort (HR 1.18, 95% CI 0.48-2.91; log-rank p = 0.72; **Supplementary Figure S4B**).

### Peripheral blood markers at diagnosis reflect tumor burden but circulating leukocyte count does not correlate with tumor immune infiltration

Routine blood parameters at diagnosis were available for 32 of 39 patients, and several correlated with aggressive disease (**Figure 7A-B**). Neuron-specific enolase (NSE), non-specific tumor marker used in NBL diagnostics, was the most consistent, being higher in INSS stage IV than I-III (median 222 vs 40 µg/l, r = 0.70), in INRG high-risk than non-high-risk disease, and in MYCN-amplified than non-amplified tumors. C reactive protein (CRP), an acute-phase marker of systemic inflammation, was higher at diagnosis in stage IV disease (r = 0.70, adjusted p = 0.047; median 26.5 vs 5.0 mg/l by stage), and age-adjusted hemoglobin was lower (r = -0.65, adjusted p = 0.047; median 94 vs 116 g/l by stage), possibly reflecting bone marrow metastases. Age-adjusted neutrophil count was also higher in high-risk disease and in tumors with unfavorable histology (r = 0.79 and 0.60), but neither association remained significant after correction for multiple testing. Lactate dehydrogenase (LD), a non-specific marker of tissue damage, was measured in 17 patients, of whom only two were stage I-III, and was therefore not formally tested here. Patients who had deceased by last-follow-up had higher neutrophil counts (median 1.9 vs 4.1 ×10□/l, p = 0.0008), CRP (5.0 vs 49.0 mg/l, p = 0.003), and NSE (63 vs 233 µg/l, p = 0.018) at diagnosis than survivors (**Figure 7C**). CRP was the only laboratory parameter prognostic in Cox regression, with higher CRP at diagnosis associated with worse overall survival (HR 7.03, 95% CI 1.90-25.97, p = 0.003).

**Figure 7.**
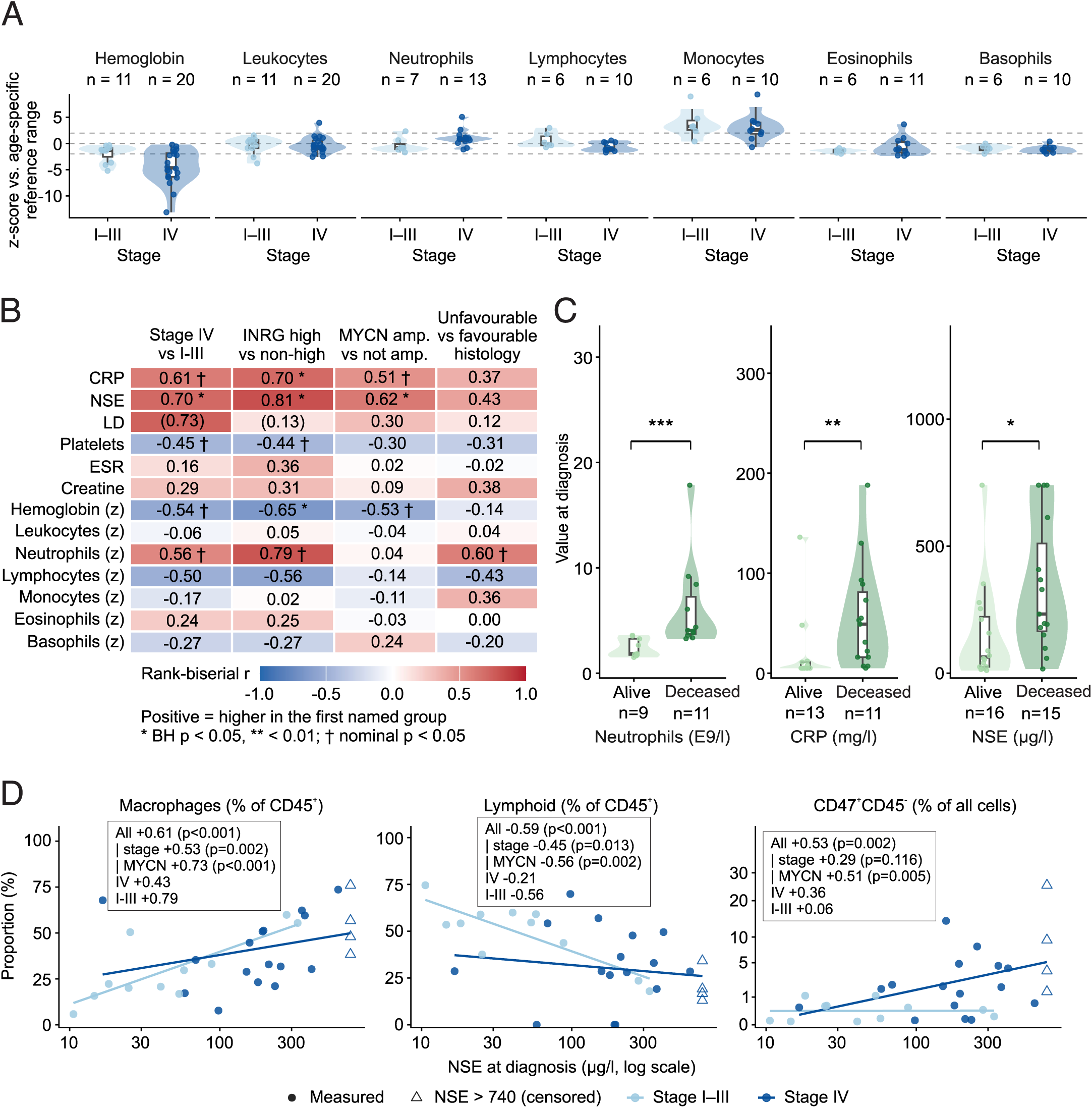
Peripheral blood markers at diagnosis are independent of tumor immune composition. **A**: Age-adjusted blood cell counts by stage. z = 0 is the middle of age-specific reference range, dotted lines its limits (z = ±1.96). Each dot is one patient; boxes show median and IQR. **B**: Laboratory values at diagnosis vs. clinical risk factors. Each cell shows the correlation from a Mann-Whitney U test. Rows marked (z) are age-adjusted z-scores calculated against pediatric reference ranges; the rest are raw values. Values in brackets = fewer than five patients per group. Benjamini-Hochberg across the grid: * adjusted p < 0.05, ** < 0.01; † nominal p < 0.05 that does not survive correction. **C**: Raw neutrophil count, CRP and NSE at diagnosis in patients alive versus deceased at last follow-up; n per panel shown in the figure. Each dot is one patient, violins the distribution, boxes median and IQR. Mann-Whitney U, * p < 0.05, ** p < 0.01, *** p < 0.001. **D**: Serum NSE at diagnosis vs. tumor microenvironment composition (n = 31; NSE not recorded for all patients). Macrophages = CD68^+^HLA-DR^+/-^CD45^+^; lymphoid = B cells (CD20^+^CD45^+^), T cells (CD3^+^CD56^−^CD45^+^), NKT cells (CD3^+^CD56^+^CD45^+^), and NK cells (CD3^−^CD56^+^CD45^+^). Each point is one patient, colored by INSS stage. Lines are least-squares fits within stage. In-panel values are Spearman correlations: across all patients, controlling for stage and for MYCN amplification, and within each stage group separately.

To assess whether blood parameters at diagnosis reflected the tumor immune infiltrate, all 14 laboratory values were correlated against 21 mIF measures (294 comparisons, BH correction within each laboratory value). Intratumoral immune infiltration was independent of the circulating leukocyte count (**Supplementary Figure S5**). Only NSE remained significant after correction for multiple testing: higher NSE correlated with a higher total TAM proportion (ρ = 0.61, adjusted p = 0.003), a lower lymphoid proportion (ρ = -0.59, adjusted p = 0.004), and a higher proportion of CD47^+^CD45^-^ cells (ρ = 0.53, adjusted p = 0.01) (**Figure 7D**). The TAM and lymphoid associations persisted after adjustment for stage. Within the stage groups, the association between NSE and TAM proportion was stronger (ρ = 0.79, p = 0.004) in stage I-III and weaker, of borderline significance, in stage IV (ρ = 0.43, p = 0.056). In contrast, the CD47^+^CD45^-^ association was largely attributable to stage: it fell to ρ = 0.29 after adjustment (p = 0.12) and was not significant within either stage group.

### The prognostic role of TIME markers

Each mIF marker was dichotomized at the cohort median and correlated to overall survival in univariate Cox models (**Table 2**). All deaths were NBL-related; overall survival therefore equals disease-specific survival in this cohort. Eleven cell populations were associated with overall survival at p < 0.05 (**Figure 8A**). Higher abundance of ten of these cell populations was associated with improved overall survival: NKT-like cells (HR 0.21, p = 0.003), PD-1^+^CD3^+^CD8^+^ T cells as a proportion of CD3^+^ cells (HR 0.21, p = 0.004, **Figure 8D**), B cells (HR 0.28, p = 0.010), CD45^+^ overall immune infiltration (HR 0.31, p = 0.018; **Figure 8B**), M2-like macrophages (HR 0.31, p = 0.018), TIM-3^+^ cytotoxic T cells (HR 0.32, p = 0.021), NK cells (HR 0.32, p = 0.023), total macrophages (HR 0.33, p = 0.024), and cytotoxic T cells as a proportion of CD3^+^ cells (HR 0.39, p = 0.048). CD47^+^CD45^-^ cells were the only marker associated with worse overall survival (HR 3.18, p = 0.024, **Figure 8E**). After adjusting for INSS stage, only high abundance of PD-1+ cytotoxic T cells remained significant for overall survival (HR 0.25, 95% CI 0.08–0.77, p = 0.015).

**Table 2.** Univariate Cox proportional hazards models for mIF markers and survival. Median split, high group compared with the low group (reference). HR > 1 = worse survival with a higher marker value. Values are patient-level medians. 19 NBL-related deaths, 22 event-free survival (EFS) events. Cox Wald p-values.

| **Marker** | **Denominator** | **OS HR<br>(95% CI)** | **p-value<br>(OS)** | **EFS HR<br>(95% CI)** | **p-value<br>(EFS)** |
|---|---|---|---|---|---|
| **Overall immune infiltration<br>(CD45<sup>+</sup>)** | total cells | 0.31 (0.12–0.82) | 0.018 | 0.58 (0.25–1.35) | 0.204 |
| **B cells<br>(CD20<sup>+</sup>CD45<sup>+</sup>)** | total cells | 0.28 (0.10–0.73) | 0.010 | 0.52 (0.22–1.21) | 0.129 |
| **T cells<br>(CD3<sup>+</sup>CD56<sup>-</sup>CD45<sup>+</sup>)** | total cells | 0.46 (0.18–1.17) | 0.102 | 0.57 (0.24–1.33) | 0.192 |
| **NKT-like cells<br>(CD3<sup>+</sup>CD56<sup>+</sup>CD45<sup>+</sup>)** | total cells | 0.21 (0.07–0.58) | 0.003 | 0.41 (0.17–0.98) | 0.044 |
| **NK cells<br>(CD3<sup>-</sup>CD56<sup>+</sup>CD45<sup>+</sup>)** | total cells | 0.32 (0.12–0.85) | 0.023 | 0.43 (0.18–1.03) | 0.058 |
| **M1-like macrophages<br>(CD68<sup>+</sup>HLA-DR<sup>+</sup>CD45<sup>+</sup>)** | total cells | 0.33 (0.12–0.86) | 0.024 | 0.59 (0.25–1.39) | 0.231 |
| **M2-like macrophages<br>(CD68<sup>+</sup>HLA-DR<sup>-</sup>CD45<sup>+</sup>)** | total cells | 0.31 (0.12–0.82) | 0.018 | 0.56 (0.24–1.31) | 0.183 |
| **Macrophages, total<br>(M1 + M2)** | total cells | 0.33 (0.12–0.86) | 0.024 | 0.59 (0.25–1.39) | 0.231 |
| **M1 share of macrophages<br>(M1/(M1+M2))** | total macrophages | 1.23 (0.50–3.03) | 0.654 | 1.00 (0.43–2.31) | 0.996 |
| **Cytotoxic T cells<br>(CD3<sup>+</sup>CD8<sup>+</sup>)** | CD3<sup>+</sup> cells | 0.39 (0.15–0.99) | 0.048 | 0.48 (0.20–1.12) | 0.089 |
| **Activated cytotoxic T cells<br>(CD3<sup>+</sup>CD8<sup>+</sup>GranzymeB<sup>+</sup>)** | CD3<sup>+</sup> cells | 0.50 (0.20–1.28) | 0.147 | 0.86 (0.37–1.99) | 0.722 |
| **Exhausted cytotoxic T cells<br>(CD3<sup>+</sup>CD8<sup>+</sup>PD-1<sup>+</sup>)** | CD3<sup>+</sup> cells | 0.21 (0.07–0.61) | 0.004 | 0.44 (0.18–1.07) | 0.069 |
| **Term. exhausted cytotoxic T cells<br>(CD3<sup>+</sup>CD8<sup>+</sup>TIM-3<sup>+</sup>)** | CD3<sup>+</sup> cells | 0.32 (0.12–0.84) | 0.021 | 0.42 (0.18–1.00) | 0.051 |
| **T helper cells (CD3<sup>+</sup>CD8<sup>-</sup> CD4<sup>+</sup>)** | CD3<sup>+</sup> cells | 0.48 (0.19–1.21) | 0.119 | 0.56 (0.24–1.33) | 0.189 |
| **Regulatory T cells<br>(CD3<sup>+</sup>CD8<sup>-</sup>CD4<sup>+</sup>FOXP3<sup>+</sup>)** | CD3<sup>+</sup> cells | 0.45 (0.18–1.16) | 0.099 | 0.79 (0.34–1.84) | 0.587 |
| **CD47<sup>+</sup>CD45<sup>-</sup>** | total cells | 3.18 (1.17–8.66) | 0.024 | 1.98 (0.83–4.76) | 0.125 |
| **CD34<sup>+</sup>CD45<sup>-</sup>Epithelial<sup>-</sup>** | total cells | 1.18 (0.48–2.91) | 0.718 | 0.90 (0.39–2.07) | 0.802 |
| **CD45<sup>+</sup>CD68<sup>+</sup>CD163<sup>+</sup>CD40<sup>+</sup> CSF1R<sup>+</sup>** | CD45<sup>+</sup>CD68<sup>+</sup> cells | 1.78 (0.70–4.53) | 0.227 | 1.47 (0.63–3.45) | 0.374 |
| **CD45<sup>+</sup>CD68<sup>+</sup>CD163<sup>+</sup>CD40<sup>+</sup> CSF1R<sup>+</sup>Clever-1<sup>+</sup>** | CD45<sup>+</sup>CD68<sup>+</sup> cells | 1.12 (0.45–2.75) | 0.811 | 0.70 (0.30–1.62) | 0.404 |
| **CD45<sup>+</sup>CD68<sup>+</sup>CD40<sup>+</sup>CSF1R<sup>+</sup>** | CD45<sup>+</sup>CD68<sup>+</sup> cells | 1.10 (0.44–2.70) | 0.843 | 0.94 (0.41–2.17) | 0.888 |
| **CD45<sup>+</sup>CD68<sup>+</sup>CD163<sup>+</sup>CD40<sup>+</sup>** | CD45<sup>+</sup>CD68<sup>+</sup> cells | 0.55 (0.22–1.38) | 0.205 | 0.62 (0.27–1.44) | 0.270 |
| **CD45<sup>+</sup>CD68<sup>+</sup>CSF1R<sup>+</sup>** | CD45<sup>+</sup>CD68<sup>+</sup> cells | 1.02 (0.41–2.54) | 0.967 | 0.87 (0.37–2.04) | 0.750 |
| **CD45<sup>+</sup>CD68<sup>+</sup>CD163<sup>+</sup>CD40<sup>+</sup> Clever-1<sup>+</sup>** | CD45<sup>+</sup>CD68<sup>+</sup> cells | 0.57 (0.22–1.51) | 0.259 | 0.76 (0.32–1.82) | 0.538 |
| **CD45<sup>+</sup>CD68<sup>+</sup>CD40<sup>+</sup>** | CD45<sup>+</sup>CD68<sup>+</sup> cells | 0.54 (0.18–1.63) | 0.271 | 0.65 (0.24–1.76) | 0.391 |
| **CD45<sup>+</sup>CD68<sup>+</sup>CD40<sup>+</sup>CSF1R<sup>+</sup> Clever-1<sup>+</sup>** | CD45<sup>+</sup>CD68<sup>+</sup> cells | 0.72 (0.27–1.89) | 0.499 | 0.73 (0.30–1.79) | 0.492 |

**Figure 8.**
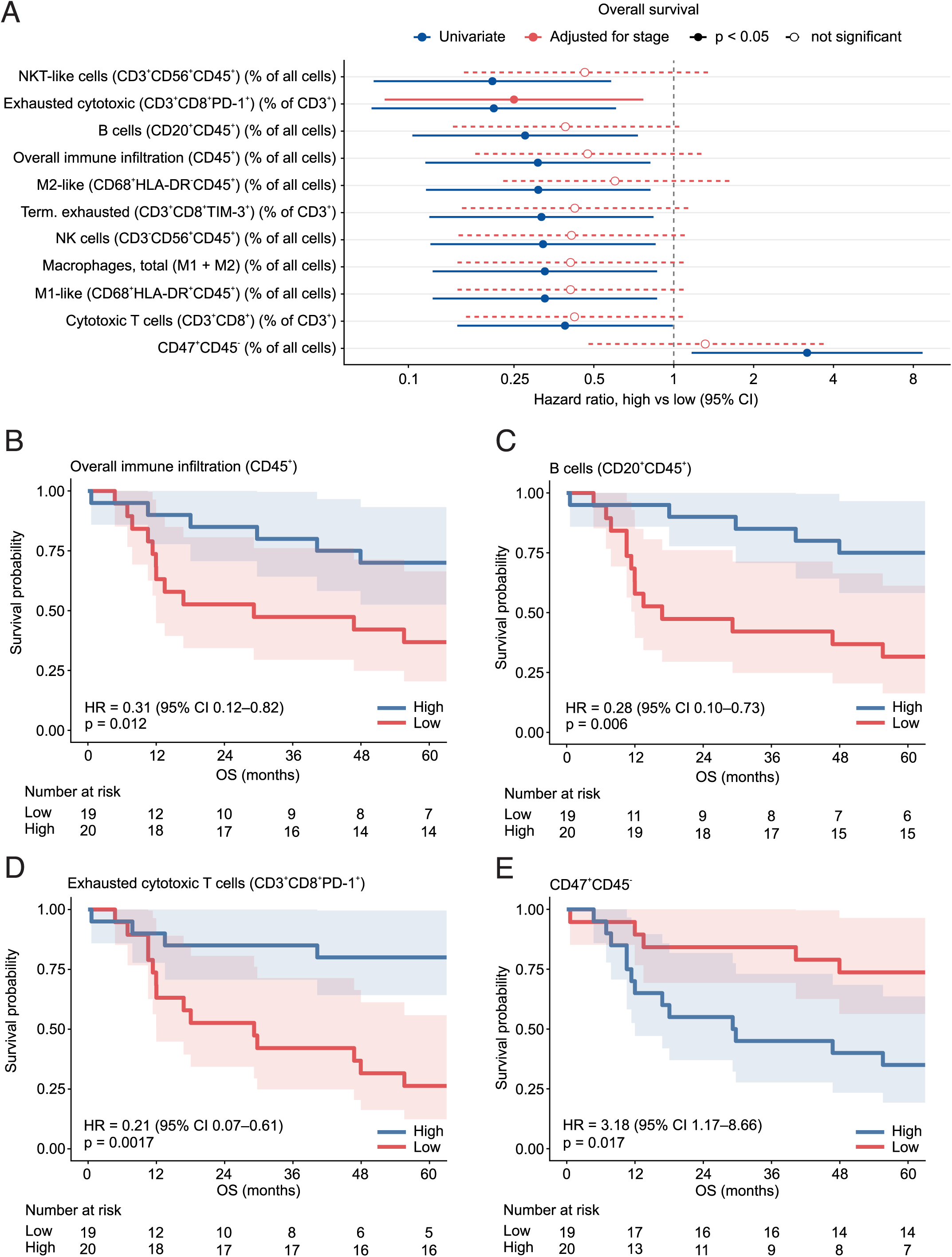
Higher immune infiltration is associated with improved overall survival in NBL. **A**: Hazard ratios for overall survival, high vs. low, for the markers reaching p < 0.05 in univariate Cox models for overall survival. Each marker is dichotomized at median; values are patient-level medians. Blue, univariate; red, adjusted for INSS stage. Solid lines with filled symbols indicate p < 0.05, dashed lines with open symbols non-significant. Overall survival in months, cutoff 5 years = 60 months. **B-E**: Kaplan-Meier curves for overall survival, median split of (**B**) overall immune infiltration (CD45^+^, % of all cells), (**C**) B cells (CD20^+^CD45^+^, % of all cells), (**D**) exhausted cytotoxic T cells (CD3^+^CD8^+^PD-1^+^, % of CD3^+^ cells) and (**E**) CD47^+^CD45^-^ cells (% of all cells). Shaded areas are 95% confidence intervals; p-values are log-rank.

No association with either endpoint was found for the stromal markers (α-SMA^+^ and PDGFRβ^+^α-SMA^+^), vascular content (CD34^+^CD45^-^Epithelial^-^), activated cytotoxic, regulatory or helper T cells, or any of the eight macrophage marker combinations (**Table 2**).

### Validation of marker-outcome associations in an external cohort

To assess whether the mIF findings carry prognostic information in an independent cohort, we analyzed bulk RNA-sequencing data from the SEQC NBL cohort (n = 498), acknowledging that bulk expression reflects tissue composition rather than cell-level protein phenotypes. In univariable Cox models with patients dichotomized at the median expression, 23 of the 28 markers shown were associated with overall survival after multiple-testing correction (**Figure 9A, Supplementary Table 1**). Higher expression predicted improved survival for the pan-leukocyte marker CD45 (*PTPRC*; HR 0.50, 95% CI 0.34-0.75), for lymphoid markers including *CD8A* (0.34, 0.23-0.52) and *CD3E* (0.47, 0.32-0.71), for myeloid markers including *CSF1R* (0.38, 0.25-0.57) and *HLA-DRA* (0.46, 0.31-0.69), and for the endothelial marker *CD34* (0.39, 0.26-0.60). The strongest associations were for CD56 (*NCAM1*; HR 0.21, 0.13-0.34) and *CD47* (0.27, 0.17-0.42). After adjustment for INSS stage, 19 markers remained significant; *CD40, CD68, TIM-3,* and *CLEVER-1* lost significance once stage was accounted for.

**Figure 9.**
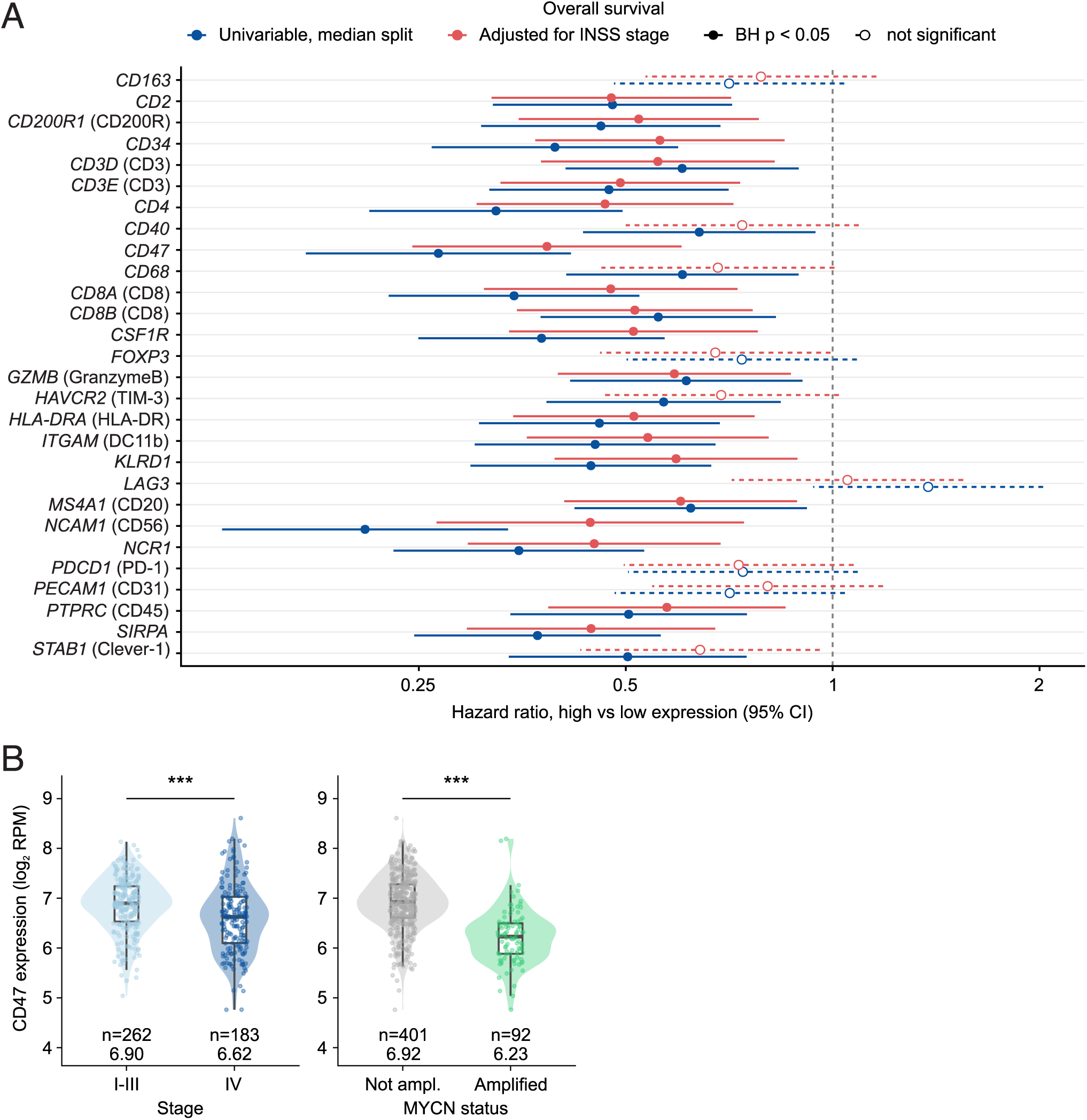
Validation of the findings in SEQC RNA dataset showed similar association between higher overall immune content and improved survival. **A**: External validation of the association of immune-related gene expression to overall survival in the SEQC NBL cohort. Hazard ratios for overall survival from Cox regression on bulk tumor RNA-seq (n = 498, 105 deaths), median split into high and low expression. Blue, univariate; red, adjusted for MYCN amplification and INSS stage IV. Bars are 95% confidence intervals. p < 0.05 was considered statistically significant. **B**: *CD47* RNA expression compared between stage I-III vs IV and MYCN amplification. *** = p < 0.001.

*CD47* remained strongly prognostic after stage adjustment (HR 0.38, 0.24-0.60), as did its receptor *SIRPa* (0.45, 0.29-0.67). Contrary to our mIF protein-level findings however, *CD47* bulk RNA expression was lower in stage IV than in stage I-III tumors (median 6.62 vs 6.90 log2 RPM, p = 7.2 × 10□□) and lower in MYCN-amplified than in non-amplified tumors (6.23 vs 6.92, p = 1.6 × 10□²²) (**Figure 9B**). The transcriptional differences were modest in magnitude, 1.2-fold by stage, and 1.6-fold by MYCN.

## DISCUSSION

NBL is a clinically heterogeneous disease, ranging from tumors with spontaneous regression to malignant treatment-refractory disease. Despite advances in risk-adapted treatment, predicting individual treatment responses remains a major challenge. In addition, patients with high-risk disease in particular require new therapeutic options such as immunotherapy. The development of these approaches depends on a detailed understanding of the TIME. In this study, we characterized the TIME of NBL using mIF staining of retrospective tissue samples from 39 patients, integrated with clinical data. Aggressive and chemotherapy-treated NBLs predominantly displayed a lymphoid-poor, macrophage-dominated TIME. In immune rich tumors, immune cell populations were spatially compartmentalized, with NK cells located closer to tumor cells than T cells. Higher immune cell infiltration was associated with improved overall survival, whereas a higher proportion of non-leukocyte CD47^+^ cells was associated with worse overall survival. TAMs frequently expressed targetable receptors including CD40, CSF1R, CD163, and Clever-1.

In our cohort, the NBL TIME was dominated by innate immune cells, whereas the adaptive compartment was sparse, consistent with previous studies ^6,13,17^. Immune infiltration was associated to disease stage: stage IV tumors had lower overall immune infiltration and lower infiltration of B cells, T cells, NK cells, and NKT-like cells. However, a large proportion of stage IV tumors in our cohort had received neoadjuvant chemotherapy prior to sampling, making it difficult to entirely separate the effects of disease stage and chemotherapy. Consistently with our findings, low cytotoxic lymphocyte, NK cell and dendritic cell content has previously been linked to high-risk, MYCN-amplified tumors and poorer prognosis ^6,9,18,19^. The low number of adaptive immune cells is consistent with the low mutational burden and low MHC class I expression characteristic of NBL ^6,20,21^. Correspondingly, neoantigen availability for T-cell priming is reported to be limited especially in high-risk disease, offering a plausible explanation for the minimal activity of T cell checkpoint monotherapy in this disease ^22^. Low MHC class I expression might make NBL cells susceptible to NK-cell recognition, which could partially explain the relative proximity of NK cells and tumor cells observed in our data.

In our cohort, the composition of the relatively small T cell compartment carried prognostic information. A high proportion of PD-1^+^ cytotoxic T cells within the CD3^+^ compartment was associated with improved overall survival independently of INSS stage, whereas the associations of total cytotoxic T cells and their TIM-3^+^ subset did not persist after adjustment for stage. Regulatory T cells were rare and not prognostic. These findings agree with previous studies linking T cell infiltration to favorable outcome in NBL ^8,19,23^. At the transcriptomic level, high-risk NBLs with a T-cell-inflamed gene expression profile have been shown to have better overall survival independent of age, MYCN status and ploidy ^22^. Although PD-1 and TIM-3 are commonly considered markers of T cell exhaustion, T cells expressing them were associated with favorable outcome in our data. PD-1 is also upregulated upon T cell activation, and PD-1^+^TIM-3^+^ co-expression develops during prolonged antigen stimulation ^24^. A high proportion of PD-1^+^CD8^+^ T cells may therefore reflect an ongoing, antigen-driven anti-tumor response rather than immune failure ^25^. In line with this, nearly all Granzyme B□CD8□ T cells co-expressed TIM-3, indicating retained cytotoxic potential within this population. PD-1 blockage monotherapy has shown minimal activity in most pediatric solid tumors including NBL ^26,27^. Our results nevertheless suggest that a subset of patients may have a pre-existing, checkpoint-restrained CD8^+^ response. However, as nearly all PD-1^+^ cytotoxic T cells co-expressed TIM-3, few progenitor-like cells are likely to remain, and PD-1 blockade alone would therefore be unlikely to restore their function. Using single-cell transcriptomics, Wienke et al. identified NECTIN2-TIGIT as a key immune checkpoint in NBL ^28^. Combination approaches, such as TIGIT and PD-1 co-blockade, or pairing checkpoint inhibition with CD47 or Clever-1 -directed therapy as discussed below, could therefore be more effective ^29^.

Induction chemotherapy has previously been shown to reshape the TIME of high-risk NBLs in multiple ways, for example by increasing total macrophage content ^12^. Several mechanisms may explain this: damage signals from dying tumor cells and the following inflammatory response attract macrophages to clear cell debris, surviving tumor cells often secrete inflammatory cytokines that attract myeloid cells, and macrophages can also proliferate locally in response to inflammation ^30,31^. Chemotherapy, cyclophosphamide in particular, induces reactive myelopoiesis which may further increase TAMs ^32^. In our study, post-neoadjuvant chemotherapy samples contained more HLA-DR^+^ macrophages while proportions of B, NKT-like and NK cells as well as HLA-DR^−^ macrophages were lower. As recently recruited, monocyte-derived macrophages typically express HLA-DR ^33^, this shift is consistent with chemotherapy-induced macrophage recruitment. Because treatment status was closely related to disease stage and risk group, some of these differences may also reflect underlying differences in tumor biology rather than treatment effects alone. However, two paired samples obtained before and after neoadjuvant chemotherapy (one stage IV tumor and one initially stage II tumor with a metastatic relapse) in our study showed changes in the same direction. As the treatment protocols have evolved over the study period, the effects of specific chemotherapy regimens could not be assessed in our cohort.

In our spatial analyses, CD56^+^CD45^−^CD31^−^PDGFRβ^−^ compartment served as a surrogate for the NBL tumor cells. CD56 (neural cell adhesion molecule, NCAM) is a glycoprotein expressed by neurons, glia, as well as NK cells and some other immune cell populations 34,35 . Although not specific to NBL, CD56 is widely expressed by NBL cells because of their neural crest origin ^36^, and even mesenchymal, GD2-negative NBL cells have been reported to retain CD56 expression ^37^. In our study, a small number of tumors were almost entirely CD56 negative, while the cores contained viable tumor tissue, as assessed by H&E staining, and the diagnosis was confirmed by a pathologist. The absence of polysialylated NCAM has been linked to unfavorable outcome in advanced NBL, but the prognostic significance of NCAM independent of its polysialylation remains unresolved ^38^. As this study focused on the immune cell compartment, other NBL markers were not included in the staining panels due to limited space. To identify tumor cells more reliably and to distinguish between adrenergic and mesenchymal-type NBL cell states, markers such as PHOX2B, HAND2, PRRX1, vimentin and CD44 should be included in addition to CD56 ^12,39–41^.

HLA-DR expression was used to generally distinguish M1-like TAMs from M2-like TAMs. HLA-DR^+^ (M1-like) TAMs dominated in most tumors. However, 90% of TAMs were also CD163^+^ and 97% CSF1R^+^, markers typically associated with M2-like, pro-tumorigenic macrophages. Thus, most TAMs simultaneously co-expressed markers associated with both M1 and M2-like phenotypes. This supports the view that the M1/M2 classification is an oversimplification of macrophage states ^42^. Previous studies have, likewise, described the variety of TAMs in NBL heterogenous, rather than organized into M1/M2 subtypes ^12,43^. HLA-DR positivity thus identifies the dominating TAM population in our samples as potential antigen-presenting cells. However, their simultaneous expression of Clever-1, which restrains antigen presentation and pro-inflammatory cytokine secretion, and the scavenger receptors CD163, and CSF1R, suggests that this presentation capacity could be functionally constrained.

Several of the macrophage receptors assessed here are being investigated as immunotherapeutic targets. Clever-1 (Stabilin-1), a scavenger receptor expressed on immunosuppressive macrophages ^44^, was present on a median of 41% of TAMs in our cohort. Interestingly, Clever-1-positive and Clever-1-negative macrophages showed consistent spatial segregation within NBL tumors. This suggests that Clever-1 expression is associated not only with macrophage phenotype but also with their spatial organization within the TIME. Whether these spatially distinct macrophage populations differ in function or in their response to Clever-1-targeted therapies remains to be investigated. In preclinical models, Clever-1 blockade has shown to reactivate cytotoxic T cell responses, and in a phase I/II MATINS trial of the anti-Clever-1 antibody bexmarilimab, high pre-treatment intratumoral Clever-1 positivity was associated with disease control ^45,46^. CSF1R and CD40 were expressed on nearly all TAMs. CSF1R signaling maintains macrophage survival and differentiation and may promote immunosuppressive TAM phenotypes, and CSF1R inhibitors have shown potential as immunotherapeutic agents ^47^. In NBL, CSF1R^+^ myeloid infiltration has been associated with poor outcome ^11^. In mice, CSF1R inhibitor BLZ945 limited NBL tumor progression ^11^. CSF1R-blockade also enhanced the efficacy of chemotherapy against NBL xenografts in T-cell depleted mice-model ^48^, which may be particularly relevant for the lymphoid-poor, high-risk tumors characterized here. CD40 receptor agonists aim to shift macrophages toward an anti-tumor phenotype and can potentiate both innate and adaptive immune responses ^49^. In an immunologically cold NBL murine model, a CD40 agonist in combination with other agents produced potent anti-tumor responses ^50^, and the CD40 agonist sotigalimab was well tolerated in a recent phase I trial in children with central nervous system tumors ^51^. The CSF1R inhibitor pexidartinib is evaluated in an ongoing phase I trial which includes pediatric solid tumors (NCT02390752) ^52^. In contrast to these targets, CD200R expressing macrophages were essentially absent in our samples, arguing against CD200R-directed approaches in NBL.

CD47 is widely expressed on normal cells and frequently overexpressed in cancer cells, and it inhibits macrophage phagocytosis by binding the receptor SIRPα ^53^. Blocking this interaction enhances macrophage clearance of tumor cells and promotes antigen cross-presentation, which in turn primes T cells for an adaptive anti-tumor response ^54^. CD47 was widely expressed in our cohort, and CD47□CD45□ cells were more abundant in stage IV and post-chemotherapy tumors. In both paired samples, the amount of CD47^+^ cells increased after chemotherapy. The proportion of CD47□CD45□ cells was the only marker associated with worse overall survival. Similarly, in osteosarcoma, high CD47 expression before chemotherapy has been associated with worse survival and its expression has been shown to increase after induction chemotherapy ^55^. Theruvath et al. showed that combining anti-GD2 dinutuximab with anti-CD47 magrolimab eradicated tumors in NBL mouse models ^16^. However, the only clinical trial of magrolimab in NBL, in combination with dinutuximab (NCT04751383), was suspended due to drug supply issues following safety concerns in adult hematological cancer trials ^56^. Given the abundance of macrophages as potential effector cells and the further increase in CD47 after chemotherapy, the CD47–SIRPα axis remains an attractive target in NBL, and next-generation agents with improved safety profiles require evaluation.

In our study, stromal content including PDGFRβ□ fibroblasts showed no association with outcome. Together with the absence of any difference in stromal content by stage, risk group, MYCN status or INPC histology, these results suggest that the stromal compartment is not a major prognostic determinant in this cohort, in line with Zeine et al. but in contrast to Hashimoto et al. ^57,58^. Tumor vascular content, estimated as the proportion of CD34^+^CD45^−^Epithelial^−^ cells, was likewise not significantly associated with stage or overall survival. Previous studies of NBL vascularity have yielded conflicting results ^59,60^. Nevertheless, the BEACON trial demonstrated that the addition of the antiangiogenic agent bevacizumab to chemotherapy improved response rates in relapsed or refractory high-risk NBL, highlighting the therapeutic relevance of tumor vasculature ^61^.

In the SEQC validation cohort, comprising bulk RNA sequencing data from 498 primary NBLs, immune gene expression was broadly associated with better overall survival, consistent with our protein-level findings. In contrast, higher *CD47* RNA expression was favorable in the SEQC dataset, whereas higher proportion of CD47□CD45□ cells was associated with worse overall survival in our cohort. This discrepancy likely reflects methodological differences. Bulk RNA sequencing captures transcripts from all cells in the sample, and CD47 is also highly expressed by red blood cells as well as endothelial, stromal and immune cells ^62^, whereas our multiplex analysis specifically quantified CD47 on non-immune (CD45□) cells and excluded red blood cells. Moreover, SEQC comprises diagnostic, mostly chemotherapy-naïve tumors, whereas the CD47 increase in our cohort was partly associated with prior chemotherapy. In the validation cohort, every significant marker-survival association in the Cox model pointed in the same direction. This might indicate that these genes largely reflect tumor composition rather than independent biological effects: aggressive NBLs are densely cellular and depleted of non-malignant cells, so bulk expression of any non-tumor gene partly measures how much non-malignant tissue is present. Consistent with this, adjustment for INSS stage attenuated the associations. Single-cell or spatial transcriptome approaches could resolve these cell-type specific effects more precisely.

To assess whether routine diagnostic laboratory markers, systemic inflammation and circulating leukocytes were associated with intratumoral immune landscape, we recorded laboratory measures at diagnosis from the patients in our cohort. Higher NSE at diagnosis associated with a macrophage-rich, lymphoid-poor TIME independently of stage. Higher CRP, NSE and neutrophils were associated with worse overall survival, as previously described ^63,64^. Circulating leukocytes did not reflect intratumoral immune infiltration, which to our knowledge has not previously been examined in NBL, but a recent study with glioblastoma demonstrated similar findings ^65^. Notably, serum NSE correlated with a macrophage-rich, lymphoid-poor TIME independently of stage, driven by HLA-DR□ (M1-like) TAMs. This association may reflect shared underlying processes, such as tumor cell damage and turnover, contributing both to NSE release in circulation and inflammatory remodeling of the TIME.

This study has several limitations. NBL is a rare disease, limiting cohort sizes in all studies; our cohort comprised only 39 NBL patients. It included samples from both lower risk (stage I-III) and high risk (stage IV) tumors but lacked stage 4S/MS tumors. Since low-risk tumors are not always biopsied, biobank material is largely enriched for higher-risk tumors. Low risk and high risk NBL are thought to be biologically very different diseases. As there were almost no deaths among stage I-III patients, the overall survival signals in the survival analysis were driven almost entirely by stage IV group, and the cohort size limited multivariable adjustment to a single covariate (stage). In this study, we used TMAs instead of whole tissue sections, meaning that larger structures and spatial immune cells organizations, such as TLSs, could not be analyzed, as previously demonstrated ^66^. Even with multiple cores per tumor, TMAs cannot fully capture intratumoral heterogeneity, particularly in tumors that responded well to chemotherapy, where viable tumor cells may be confined to small areas. Both TMAs and whole-slide sections are also two-dimensional samples of three-dimensional tissue, so spatial relationships may differ in other planes. Finally, the limited number of markers per panel required staining separate sections, which prevented cell-level integration across panels and the inclusion of multiple tumor cell markers for robust identification of NBL cells.

## Conclusions

High-risk and chemotherapy-exposed NBL tumors showed a lymphoid-poor, macrophage-rich immune microenvironment with prominent expression of potential macrophage-related immunotherapy targets, and distinct spatial organization of immune cell populations. Elevated serum NSE was associated with a similar immune phenotype independently of stage, potentially reflecting shared processes of tumor-cell damage and inflammatory remodeling. Together, these findings highlight the potential clinical relevance of immune composition and spatial organization in NBL, suggest that routinely measured serum NSE may provide insights into tumor immune biology, and support further investigation of macrophage-directed therapeutic strategies. Further studies with a larger patient cohort are needed to confirm these findings.

## METHODS

### Patient cohort and clinical data

Formalin-fixed paraffin-embedded (FFPE) NBL tissue samples were obtained from Helsinki Biobank at Helsinki University Hospital. All available NBL samples were requested, and samples with sufficient FFPE tissue for tissue microarray (TMA) building were included in the cohort. Samples were originally collected at the time of biopsy or tumor resection at Children’s Hospital, Helsinki University Hospital between 1995-2018 and informed consent was obtained when samples were deposited to the Helsinki Biobank. The study was approved by Helsinki University Hospital Regional Committee on Medical Research Ethics (HUS/2674/2021), conducted in accordance with Finnish bylaws, and adhered to the principles of Declaration of Helsinki.

Clinical data were collected from electronic patient records. The date of radiological diagnosis was defined as the date when tumor was first noted on any type of imaging. Chemotherapy-naïve tissue was available for 15/39 patients. Neoadjuvant chemotherapy regimen varied over the years (**Supplementary Table 2**). Needle biopsy material was excluded as it could not be utilized for the TMA. Tumors were staged according to the International Neuroblastoma Staging System (INSS), as this was the only staging system recorded at diagnosis for most patients, and image-defined risk factors were not systematically documented in older cases. Patients were assigned to International Neuroblastoma Risk Group (INRG) risk groups based on the clinical data available at data collection. Patients with insufficient data, such as missing MYCN or chromosomal status, were classified as unknown. MYCN status at the time of diagnosis was recorded if available. High amplification was defined as >8x FISH (4-fold increase). Values over 2 but under high amplification criteria were marked as low amplification. Laboratory values closest to radiological diagnosis date, +/- 3 days, were recorded. Only values before the start of treatment were noted. Age-dependent blood counts were expressed as z-scores against age-specific pediatric reference ranges (**Supplementary Table 3**). Cut-off for follow-up was 31^st^ December 2024. Patients were numbered NBL01-NBL40. NBL18 was later excluded because the sample contained no viable tumor tissue, leaving 39 patients for analysis.

### Construction of tissue microarrays

The construction of TMAs was done by Helsinki Biobank (Helsinki, Finland). Fresh H&E-stained whole slide images were prepared from the selected FFPE tissue blocks and digitized using a Pannoramic 250 Flash III microscope slide scanner (3DHISTECH Ltd., Budapest, Hungary) in bright-field mode with a 20x objective. Pathologist annotated 1mm cores containing viable tumor from the digitalized whole-slide H&E images using SlideViewer v.2.6 software (3DHISTECH). For each tumor sample, 4 replicate cores from the tumor center and 1 core from adjacent non-cancerous tissue (if available) were annotated. In addition, a core from the tumor border was annotated if possible. Altogether, 185 TMA cores were collected (1-9 per patient, 1-6 per sample). Of these, 158 were from tumor center, 23 from non-cancerous adjacent tissue and 4 from tumor border. The number of cores per sample varied since some samples did not contain enough tissue material to obtain the full set of cores. The cores were mostly collected from one sample per patient; however, two patients (NBL23 and NBL29) contributed with a longitudinal sample pair before and after neoadjuvant chemotherapeutic treatment. The layout of the two TMA blocks was designed, and digitally annotated images were overlaid with donor block images. TMA blocks were constructed using the TMA Grand Master tissue microarrayer (3DHISTECH).

### Multiplexed immunofluorescence staining

In-house multiplexed immunofluorescence (mIF) staining protocol was used to stain FFPE TMA microarrays ^67^. TMA blocks were sectioned at a thickness of 4μm using a rotary microtome. Sections were mounted onto positively charged glass slides and air-dried at room temperature (RT) overnight to ensure proper adhesion. Prior to mIF staining, slides were deparaffinized in an oven at 60 °C and ran through paraffin removal program with using the Sakura Tissue-Tek Prisma Plus Automated Stainer (Sakura Finetek, Torrance, CA, USA). Epitopes in deparaffinized tissue sections were made available for binding through heat induced epitope retrieval (HIER) (immersion in hot (99□°C) EDTA-based pH 9 buffer for 20□min) (PT Module, Thermo Fisher, USA). Endogenous peroxide activity was blocked in 0.9 % H_2_O_2_ for 15□min at RT, followed by protein blocking in 10% normal goat serum (NGS) in TBS for 30□min at RT. Primary antibodies were diluted to 10% NGS-TBS solution and incubated on slides for 1.5h at RT.

Primary antibodies and their information are listed in **Supplementary Table 4**, marker panels and staining rounds are shown in **Table 3**. First round of staining was done by incubating the slides with the primary antibodies for 2h after which they were washed with PBS and secondary anti rabbit horseradish peroxidase conjugates were incubated with the samples for 30 min after which tyramide signal amplification (TSA) fluorophore was applied for 15 min. The slides were then washed and the goat anti-mouse HRP conjugated antibody then applied, and second TSA fluorophore applied. Antibodies were then deactivated by HIER procedure and second round of primary antibodies applied. After an hour incubation the secondary Alexa Fluor fluorophore conjugated goat anti mouse and anti-rabbit antibodies were applied together with DAPI (secondaries and DAPI diluted in 1:300 TBST) for 30 min, after which the slides were dried, cover slipped and then imaged. Between the staining rounds the previous fluorophores were bleached using a 4.5% H_2_O_2_ 0.024 M NaOH-TBS solution under direct light and the previous antibodies denatured using the HIER procedure. For second to fifth round of stains, the primary antibodies were applied for 1 h after which Alexa Fluor fluorophore conjugated goat antibodies (1:300 in TBST) were applied together with 1:150 diluted DAPI for 30 min. The slides were cover slipped after this step.

**Table 3.** Multiplexed immunofluorescence panels.

| **Round** | **Channel** | **Panel 1:**<br>**TME** | **Panel 2:**<br>**T cells** | **Panel 3: CAF** | **Panel 4:**<br>**IOTT** |
|---|---|---|---|---|---|
| **1** | TSA-488 | CD68 | PD-1/CD279 | PDGRFβ | CSF1R |
| **1** | TSA-555 | CD3 | CD3 | FAP | CD47/IAP |
| **1** | Alexa-647 | CD56/NCAM | PD-L1 | pSTAT3 | MYCN |
| **1** | Alexa-750 | CD11b/ITCAM | CD8 | SMA | CD200R |
| **2** | Alexa-647 | PDGRFβ | TIM3 | Collagen I | CD45 |
| **2** | Alexa-750 | CD20 | FOXP3 | PanCK + E-cadherin | CD68 |
| **3** | Alexa-647 | CD31/PEKAM1 | Granzyme B |  | CD34 |
| **3** | Alexa-750 | CD45 | Ki-67 |  | WT1 |
| **4** | Alexa-647 | HLA-DR | CD4 |  | CLEVER1/STAB1 |
| **4** | Alexa-750 | PanCK + E-cadherin | PanCK + E-cadherin |  | CD163 |
| **5** | Alexa-647 |  |  |  | CD40 |
| **5** | Alexa-750 |  |  |  | PanCK + E-cadherin |

The mIF slides were imaged using Axio Scan.Z1 (Zeiss, Germany) with a 20x objective (NA 0.8) at the Digital Pathology (DP) core unit (Institute for Molecular Medicine Finland-FIMM, HiLIFE, University of Helsinki, Finland).

### Cell-based image analysis

Prior to downstream analysis, a multi-step pre-processing workflow was applied to the mIF images. First, low-resolution (10%) TIFF images containing only the DAPI channel were generated using Zen Light (Zeiss, version 3.1). These images were used to annotate individual TMA cores in QuPath ^68^, enabling precise identification and localisation of each core within the full TMA slide. Based on the generated annotations, single-core TMA images were extracted from the original whole-slide images using a custom Python pipeline. This step produced individual TMA core images at both full resolution and reduced resolution (25% TIFFs), facilitating efficient downstream processing.

To remove autofluorescence from red blood cells (RBC) we generated a mask identifying them for exclusion. The 25% resolution TMA images from the first staining round were converted into RGB images using CellProfiler ^69^. These RGB images were subsequently imported into ilastik (version 1.4.0) ^70^, where supervised pixel-level classification was performed to annotate red blood cells, tissue regions, and empty background. The resulting probability maps were used to create RBC masks that were applied in later analysis steps to exclude non-tissue regions and correct for blood-related signal artefacts.

The image registration and cellular segmentation followed in-house Python pipeline (https://github.com/bioimage-profiling/iCAN-tma-general-profiling). Shortly, nuclear DAPI signals from the different imaging rounds were aligned by finding optimal translation transformation based on phase cross-correlation. The optimal parameters of each round were used to transform all channels of that round.

Cellular segmentation was performed by first segmenting the nuclear regions in DAPI channel images using the cellpose (v.3.0.7) pretrained *nuclei* model followed by filtering out smaller than 50px sized nuclei ^71^. Cell regions were defined by dilating the nuclear segmentation by a disk-shaped structuring element with a 3px radius. All cells overlapping more than 25% with the autofluorescence mask were removed from further analysis. Total cell numbers yielded per patient are shown in **Supplementary Table 5**. We set cell mean intensity thresholds separately for each channel to define whether a cell was marked positive or negative for a marker. Thresholds were set separately for each glass of TMAs to reduce possible bias due to sample preparation and batch effects.

### Spatial analyses

Spatial analyses were performed using single-cell x–y centroid coordinates from individual TMA cores. Distances were calculated in image-coordinate space and are reported as relative observed-to-expected ratios rather than absolute physical distances. For analyses involving multiple immune-cell populations simultaneously, cells were assigned to mutually exclusive phenotypes using the CD68-prioritized hierarchical classification. M1-like macrophages were defined as CD45^+^CD68^+^HLA-DR^+^ and M2-like macrophages as CD45^+^CD68^+^HLA-DR^−^. Among CD68^−^ immune cells, B cells were assigned first as CD45^+^CD20^+^ cells. The remaining CD20^−^ cells were classified as NK cells (CD45^+^CD3^−^CD56^+^), T cells (CD45^+^CD3^+^CD56^−^), or NKT-like cells (CD45^+^CD3^+^CD56^+^). Thus, CD68 and CD20 exclusion criteria were applied in these multi-population spatial analyses to ensure mutually exclusive phenotype assignment, while the underlying predefined immune-cell phenotypes remained unchanged. For the focused B–NK analysis, the predefined B-cell (CD45^+^CD20^+^) and NK-cell (CD45^+^CD3^−^CD56^+^) phenotypes were used directly. Immune-rich tumors were defined as the upper quartile of patient-level CD45^+^ immune-cell infiltration, with 10 of 39 patients contributing 36 malignant tumor cores. Pairwise immune-cell spatial organization was assessed using bidirectional cross-nearest-neighbor distances. For each pair, the observed distance was compared with a null distribution generated by 999 random permutations of phenotype labels among the pooled cells of the two populations within the same core, while preserving cell coordinates and the number of cells assigned to each phenotype. A core was considered evaluable when at least 10 cells of each phenotype were present. Observed-to-random ratios >1 indicated greater spatial separation than expected under random labeling, whereas ratios <1 indicated relative spatial proximity. For patient-level analyses, the median observed-to-random ratio across evaluable tumor cores was calculated for each patient, and deviation from a ratio of 1 was assessed using a Wilcoxon signed-rank test.

To evaluate whether spatial segregation extended across the broader immune compartment, all 15 pairwise combinations of the six mutually exclusive immune phenotypes were analyzed using the same random-label framework. Patient-level median observed-to-random ratios were calculated across evaluable cores. For visualization, these ratios were log2-transformed, such that positive values indicated spatial separation, values near zero indicated the random-label expectation, and negative values indicated relative proximity. Patient-level ratios were compared with 1 using Wilcoxon signed-rank tests, with Benjamini–Hochberg correction across the 15 phenotype pairs. Comparisons represented by fewer than eight evaluable patients or fewer than 15 evaluable tumor cores were considered to have limited coverage and were interpreted as exploratory.

Localization of immune-cell populations relative to the CD56^+^CD45^−^CD31^−^PDGFRβ^−^ tumor-associated compartment (hereafter called CD56^+^ tumor cells) was analyzed across the cohort. CD56^+^ tumor cell positions were held fixed. For each classified immune cell, the distance to the nearest CD56^+^ tumor cell was calculated. For each immune phenotype within a tumor core, the observed mean distance to CD56^+^ tumor cell was divided by the mean CD56^+^ tumor cell distance of all six pooled classified immune-cell populations in that core. The pooled immune-cell mean represents the expected phenotype distance under random reassignment of immune phenotype labels among the fixed immune-cell positions. Thus, ratios <1 indicate relative localization closer to a CD56^+^ tumor cell than the pooled immune-cell distribution, whereas ratios >1 indicate relatively more distal localization. Cores were considered evaluable when they contained at least 10 CD56^+^ tumor cells and at least 10 cells of the immune phenotype being analyzed. Patient-level values were calculated as the median ratio across evaluable tumor cores. Wilcoxon signed-rank tests against a ratio of 1 were performed for each phenotype, with Benjamini–Hochberg correction across the six immune phenotypes.

Spatial maps were generated directly from single-cell coordinates and by using the same immune-cell classification and CD56^+^ tumor cell definition as in the quantitative analyses. Representative cores were selected to illustrate spatial patterns observed in the quantitative analyses.

### Descriptive analysis of tumor-border and tumor-adjacent non-cancerous tissue

To explore immune-cell changes across tissue regions, tumor cores, adjacent non-cancerous tissue cores, and tumor-border cores were compared descriptively. Tumor-border tissue was available from four patients (NBL09, NBL25, NBL33, and NBL37). For B cells, NK cells, and M2-like macrophages, phenotype representation was expressed as a proportion of CD45^+^ cells. Where multiple cores of the same tissue region were available from a patient, the patient-level median was used. Because only four patients had tumor-border tissue available, no inferential statistical testing was performed for these comparisons.

For visualization, patients with matched tumor and adjacent non-cancerous tissue were shown as background trajectories, while the four patients with an additional tumor-border core were highlighted across the tumor–tumor-border–adjacent tissue sequence. NBL33 was selected as a representative spatial example because its tumor-border core showed the clearest histological tumor–border interface. Spatial maps of the NBL33 tumor core, tumor-border core, and adjacent non-cancerous tissue were generated using the same single-cell phenotype definitions and Gate C classification described above.

### RNA sequencing data analysis

SEQC dataset consisting of 498 NBL patients was obtained from Gene Expression Omnibus (accession number GSE62564 for RNA sequencing data and GSE49711 for clinical information, retrieved 4 October 2026). Data contains log2 RNA expression values using RPM normalization as well as information on clinical variables including stage, MYCN status and overall and event-free survival. The cohort includes 105 deaths and 183 events; 92 MYCN amplified; INSS stage 1/2/3/4/4S = 121/78/63/183/53.

Three Cox proportional hazards models were fitted for each gene and each endpoint: univariable with expression as a continuous log2 variable, univariable with patients dichotomized at the cohort median into high and low expression, and the median split adjusted for INSS stage 4 (stage 4S grouped with the non-stage-4 patients). Hazard ratios are reported with 95% confidence intervals and Wald test p-values. P-values were corrected for multiple testing using the Benjamini-Hochberg method across the 52 genes, separately within each model and endpoint, and BH-adjusted p < 0.05 was considered significant.

### Statistical analyses

Statistical tests are indicated in each figure legend. In general, p values < 0.05 were considered significant. Statistical analyses were performed in Rstudio (R version 4.6.0) using survival, survminer and ggplot2 libraries and the tidyverse packages or in Jamovi (version 2.6.44). Normality of the data was estimated by using Shapiro Wilks p-value. When the data was not normally distributed, Mann-Whitney U test was used. For correlation analyses, two-sided Spearman rank correlations were used on patient-level medians. When multiple correlations were tested together, Benjamini-Hochberg adjusted p-values were used. For survival analyses from own data, hazard ratios were calculated using Cox proportional hazards model. Overall survival (OS) curves were created using Kaplan-Meier and compared by log-rank test. 95% confidence interval is shown with the curves. The median value of tumor cores per patient was used. NBL23 and NBL29 had paired samples and chemotherapy-naïve sample values were used in the survival analysis.

## AUTHOR CONTRIBUTIONS

Conceptualization: R.N., M.Po., K.E., V.Pi. Methodology: R.N., M.Po., S.L., M.K., V.Pe., K.E., V.Pi, L.P. Investigation: R.N., M.Po, L.P, V.V., A.M. Writing – Original Draft: R.N., M.Po., M.Pi., K.E., V.Pi. Writing – Review & Editing: R.N., M.Po., L.P., A.M., V.V., S.L., M.K., V.Pe., M.Pi., K.e., V.Pi. Visualization: R.N., M.Po., M.Pi. Funding Acquisition: V.Pi., K.E., M.Po. Supervision: V.Pi., K.E., M.Pi.

## Supporting information

All Supplementary Figures

## ACKNOWLEDGEMENTS

This study was supported by funding from the Cancer Society of Finland, HUS Helsinki University Hospital, the Helsinki Institute for Life sciences (Proof of concept grant), the Research Council of Finland, iCAN Digital Precision Cancer Medicine Flagship (iCAN), the Foundation for Pediatric Research, the Mary and Georg C. Ehrnrooth Foundation, the Väre Foundation, the Aamu Foundation, the Päivikki and Sakari Sohlberg Foundation, and the Kymenlaakso Regional Fund of the Finnish Cultural Foundation. The authors wish to acknowledge CSC – IT Center for Science, Finland, for computational resources. The samples used in the analyses were obtained from the Helsinki Biobank. We thank all study participants for their generous participation in the Helsinki Biobank. We thank Emil Eriksson for his contribution to the image analysis work. We gratefully acknowledge the FIMM Digital Microscopy and Molecular Pathology Unit (HiLIFE – Helsinki Institute of Life Science, University of Helsinki, Finland).

## DATA AND CODE AVAILABILITY

The mIF data is currently available upon request. Information for code used are listed in the Methods sections.

