## Supplementary material for "Neuroblastoma immune microenvironment is characterized by limited T cell infiltration and abundant macrophages expressing targetable receptors": All Supplementary Figures

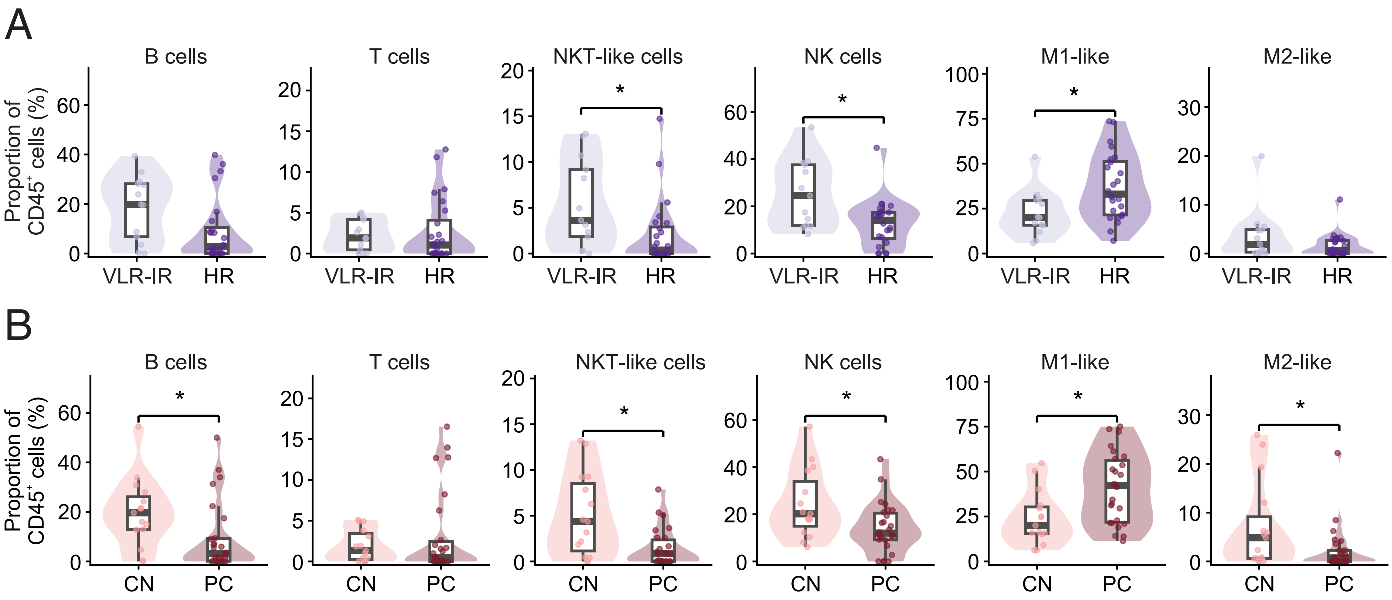


**Supplementary Figure S1. Overall composition of neuroblastoma tumor immune microenvironment.** Comparison of main immune cell populations between INRG risk groups: very low, low and intermediate risk (VLR, LR, IR; n = 13) vs high risk (HR; n = 24); two patients with unknown risk group excluded. (**A**)

Comparison of chemotherapy-naive samples (chemotherapy-naïve, CN; n = 15) and samples treated with neoadjuvant chemotherapy prior to sample extraction (post-chemo, PC; n = 26), two patients with paired samples are shown in both groups. Chemotherapeutics regimen varied between samples. (**B**)

Each dot represents the median of tumor cores from each patient. Boxplot shows the median and quartile values with whiskers extending to the most extreme non-outlier values (1.5 × IQR). P-values were calculated using Mann-Whitney U test with Benjamini-Hochberg FDR correction. * = p-value < 0.05.

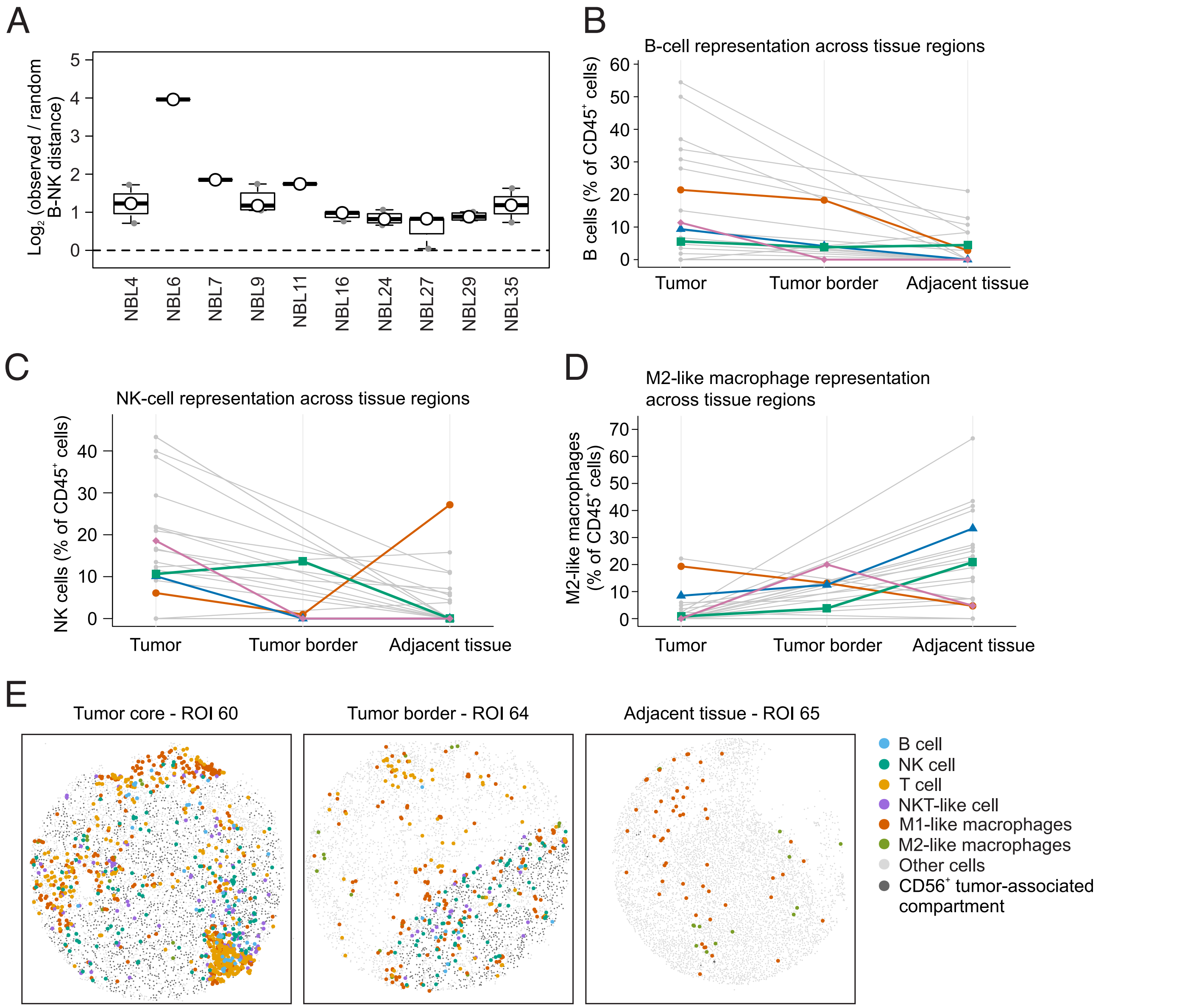


**Supplementary Figure S2. Exploratory spatial and tissue-region patterns of B cells, NK cells, and M2-like macrophages in neuroblastoma.**
Spatial separation between B and NK cells in immune-rich tumor cores. Cores containing at least 10 B cells and at least 10 NK cells were considered evaluable. Spatial separation was quantified as the observed B–NK nearest-neighbor distance relative to the expected distance after random reassignment of B- and NK-cell labels within each core; values >1 indicate greater-than-expected separation. (**A**)

Patient-level representation of B cells (**B**), NK cells (**C**), and M2-like macrophages (**D**) across tumor, tumor-border, and adjacent non-cancerous tissue regions. Values represent median phenotype proportions among CD45+ cells within each tissue region. Gray lines indicate other patients with matched tumor and adjacent tissue, whereas colored lines indicate the four patients with an additional tumor-border core (NBL09, NBL25, NBL33, and NBL37). Tumor-border comparisons were considered descriptive because only four patients had available border tissue.

Representative spatial maps from NBL33 showing a tumor core (ROI60), tumor-border core (ROI64), and adjacent non-cancerous tissue. Maps show mutually exclusive immune-cell phenotypes together with the CD56+CD45−CD31−PDGFRβ− tumor-associated compartment. Immune phenotypes were assigned using the CD68-prioritized hierarchy used in the spatial analyses. (**E**)

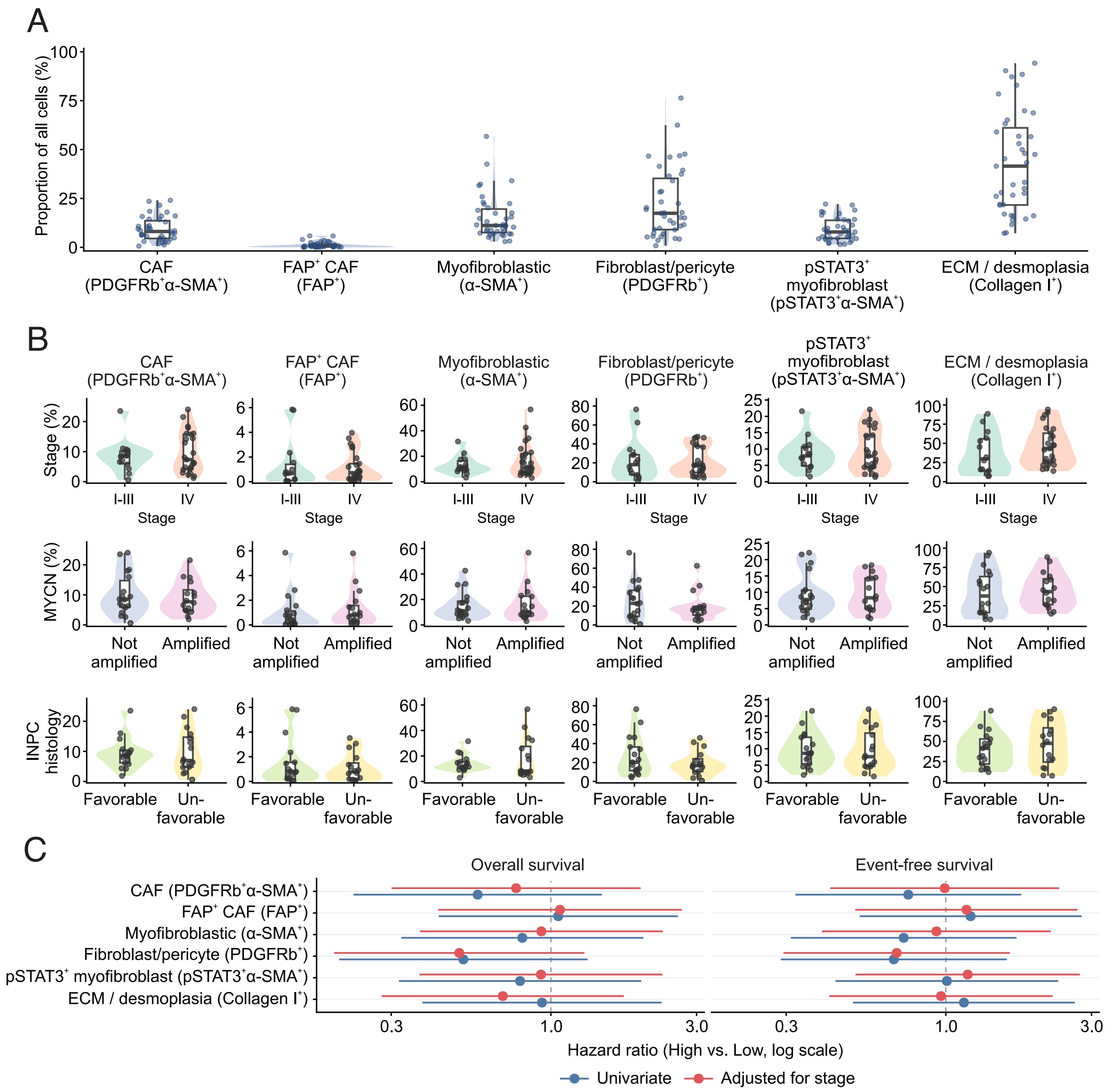


**Supplemetary Figure S3. Stromal and cancer-associated fibroblast markers in neuroblastoma.** Stromal and cancer-associated fibroblast (CAF) marker composition in NBL. Expression of six stromal markers shown as a proportion of all cells. (**A**) All populations are gated Epithelial-negative. Each dot represents the median of one patient.

Stromal and CAF marker expression in different NBL subgroups (**B**). INSS stage I–III (n = 14) vs IV (n = 25); MYCN not amplified (n = 18) vs amplified (n = 16); favorable (n = 17) vs unfavorable (n = 16) INPC histology. Patients with unknown MYCN status (n = 5) or histology (n = 6) were excluded from the respective rows. Mann-Whitney U with BH correction within each comparison; none reached significance.

Stromal content and survival (**C**). Hazard ratios for high vs low stromal content, median split, for overall and event-free survival (n = 39; 19 deaths, 22 event-free survival events). Blue = univariate Cox model, red = adjusted for INSS stage. BH correction was applied. No stromal measure reached significance. Values are patient-level medians.

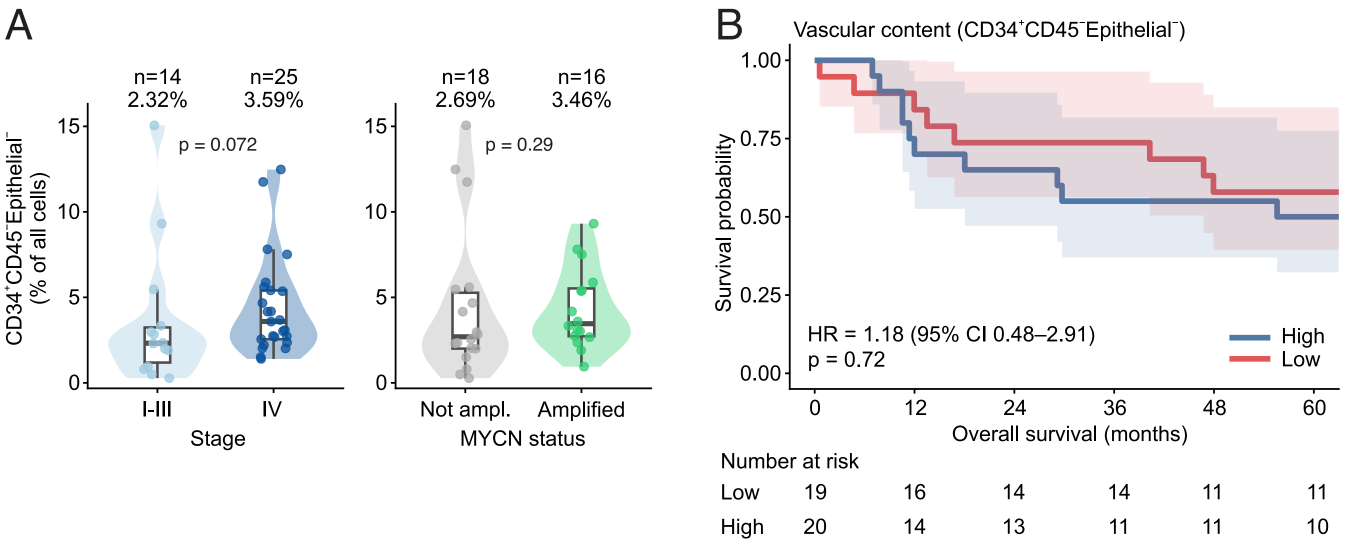


**Supplementary Figure S4. CD34⁺ vascular content in neuroblastoma tumors.**

$CD34^+CD45^-Epithelial^-$ cells, a surrogate for vascular content, as a proportion of all cells, compared between INSS stage I-III and stage IV tumors and between MYCN non-amplified and amplified tumors (**A**). Each dot is one patient (n = 39), values are medians for each patient, with boxes showing median and interquartile range. Mann-Whitney U for p-value, neither reached significance.

Kaplan-Meier curve for overall survival with patients split at the median of $CD34^+CD45^-Epithelial^-$ cell content (low, n = 19; high, n = 20; **B**). Shaded areas are 95% CI, HR is for high versus low expression; log-rank p-value shown.

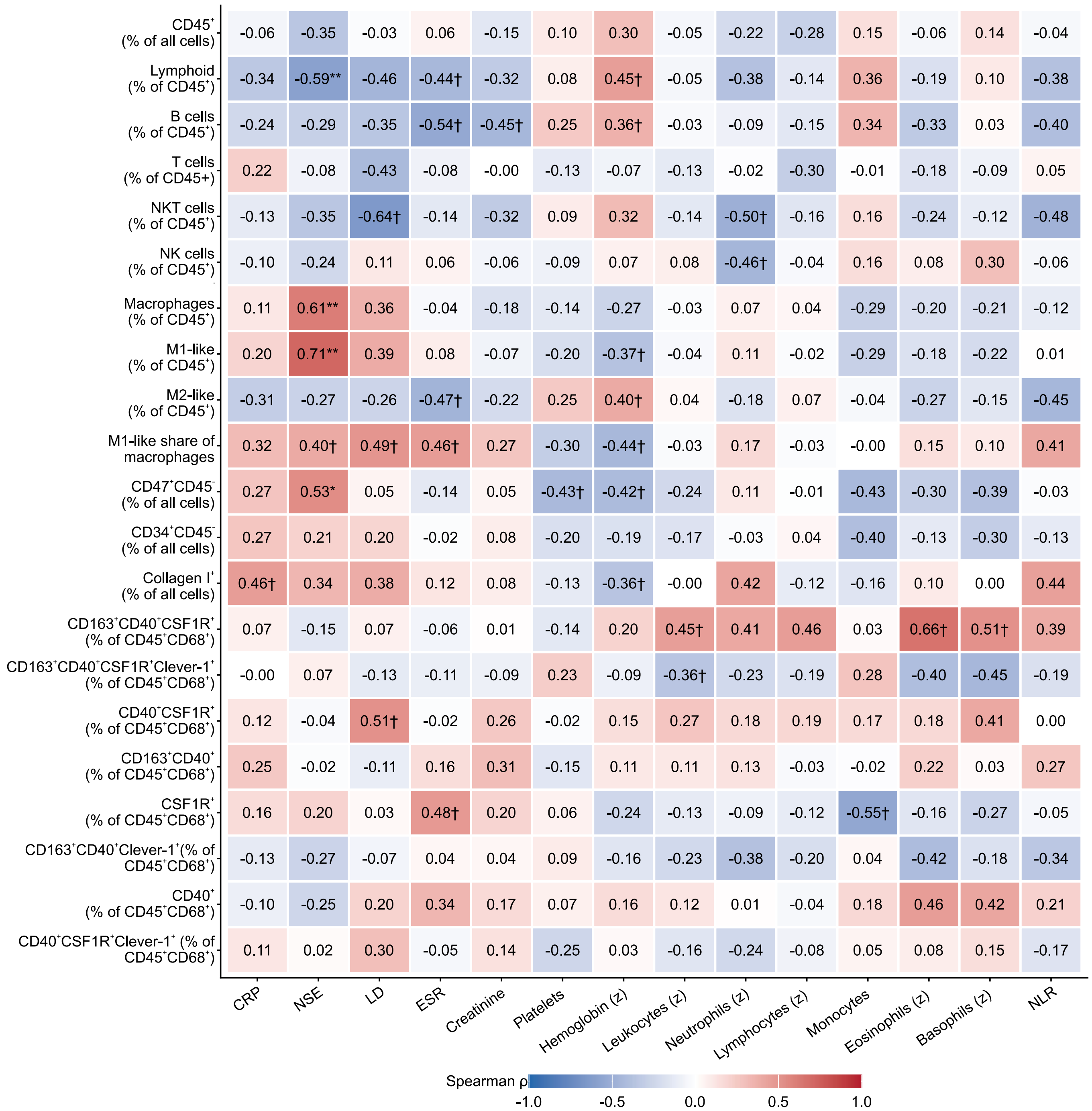


Exploratory. Benjamini-Hochberg correction within each laboratory marker (column).
* adjusted $p < 0.05$, ** $< 0.01$; † nominal $p < 0.05$ only; n.t. = fewer than 6 pairs.

**Supplementary Figure S5. Correlations between blood values at diagnosis and tumor microenvironment markers.**

Correlations between blood values at diagnosis and tumor microenvironment composition. Heatmap of Spearman correlation coefficients for all 14 laboratory markers (columns) against 22 mIF measures (rows); the coefficient is printed in each cell and encoded by color. Values are per patient: laboratory values measured at diagnosis and mIF measures the median across each patient's malignant tumor cores. The number of patients varies between cells (n = 16-39) because of not available laboratory values. Benjamini-Hochberg correction was applied within each laboratory marker across the mIF measures: * adjusted $p < 0.05$, ** $< 0.01$; † nominal $p < 0.05$ that does not survive correction.
